# Distinct amygdala-striatal and amygdala-cortical circuits encode and retrieve predictive learning to guide choice and decision-making

**DOI:** 10.64898/2026.10.12.758667

**Authors:** Elise Pepin, Billy C. Chieng, Nura Lingawi, Beatrice K. Leung, Vincent Laurent, Bernard W. Balleine

## Abstract

It has long been known that Pavlovian cues can influence choice between instrumental actions, although how this is achieved at a neural level is not fully understood. Here we examined the neural processes engaged during the encoding and retrieval of the Pavlovian stimulus-outcome (S-O) associations prior to and during their influence on choice. Using a Pavlovian-instrumental transfer paradigm (PIT), we found, as previously reported, that a circuit connecting the basolateral amygdala (BLA) and nucleus accumbens shell (BLA→NAc-S) is necessary to encode specific S-O associations within the NAc-S. However, this circuit was not required for the retrieval of those associations: optogenetic inhibition of this circuit had no effect on the ability of specific S-O associations to influence choice between instrumental actions. The BLA also projects to the infralimbic cortex (IL) although, using circuit mapping, we found that this projection was largely independent of the projection to the NAc-S. Nevertheless, using a circuit-specific disconnection approach, we found that the BLA→IL circuit is necessary for both the encoding and retrieval of specific S-O associations to influence choice. Furthermore, expanding on this disconnection approach, optogenetic inhibition of the BLA→IL circuit in one hemisphere and the IL→NAc-S contralaterally provided evidence that the broader BLA→IL→NAc-S circuit is necessary to retrieve specific S-O associations to influence choice during PIT. These data reveal, therefore, evidence of an amygdala-cortico-striatal circuit necessary for the retrieval of predictive learning to influence choice that differs from the circuit involved in encoding those predictive associations.

## INTRODUCTION

The ability of predictive learning between environmental stimuli and motivationally important outcomes to influence choice between instrumental actions is a fundamental feature of adaptive behavior (1). These predictive associations are, however, often encoded well before they are retrieved during choice, requiring the formation of enduring stimulus-outcome (S-O) memories to influence future actions. The outcome specific Pavlovian-instrumental transfer (PIT) paradigm provides a powerful experimental framework for studying this effect of predictive learning. In this task, Pavlovian stimuli predicting distinct outcomes are presented while subjects choose between instrumental actions earning those same outcomes and, in both humans and rodents, such stimuli have been found to bias choice towards the action earning the outcome they predict (2–4). PIT offers, therefore, both a behavioral measure of the influence of S-O associations on choice and a tool with which to identify the neural bases of that influence.

Regarding the latter, a substantial body of evidence suggests that PIT depends on a circuit involving the basolateral amygdala (BLA) and nucleus accumbens shell (NAc-S). The BLA is critical for encoding and retrieving sensory-specific S-O associations, and disrupting BLA function abolishes the ability of reward-predictive stimuli to influence choice during PIT (5–10). By contrast, the NAc-S appears to play little role in the acquisition of either Pavlovian or instrumental learning yet is essential for the expression of specific PIT: rats with NAc-S lesions acquire Pavlovian conditioned responding and instrumental performance normally but fail to show transfer (7, 11–14). Early findings suggested that a BLA*→*NAc-S circuit is necessary for Pavlovian learning to influence instrumental performance: disconnecting the two structures using asymmetrical lesions abolished PIT (8). More recent evidence has established that the direct BLA*→*NAc-S projection is essential for encoding specific S-O associations that support transfer. Within the NAc-S, converging neurotransmitter and neuromodulatory signals, including dopamine, glutamate, acetylcholine and opioids, interact to regulate incoming information and support the integration necessary for stimulus-based choice (7, 15, 16). The mechanisms by which this occurs at the level of molecules and circuits are complex; nevertheless, work from our laboratory has highlighted one particularly compelling example of anticipatory plasticity involving cholinergic interneurons (CINs) in the NAc-S (7, 13, 14, 17). Considerable evidence suggests that, during Pavlovian conditioning, CINs undergo a form of learning-induced plasticity in which exposure to S-O contingencies causes the accumulation of G protein-coupled δ-opioid receptors (DOPrs) on their somatic membrane.

Importantly, DOPr accumulation depends on the BLA: inhibiting (15) or exciting (7) the BLA during Pavlovian conditioning blocks or enhances DOPr expression on CINs, respectively. A defining feature of this plasticity is its prospective nature. As is true of NAc-S function generally, DOPr signaling tracks both the strength of the Pavlovian conditioned response and the Pavlovian contingency (17) but is not necessary for Pavlovian conditioning per se. It is, however, necessary for PIT (7) and, as such, appears to prepare the NAc-S circuitry for future behavioral demands, rendering specific S-O associations available to influence subsequent choice between actions. Consistent with this account, deleting CINs or blocking their function in the NAc-S, deleting DOPrs from CINs, or preventing the BLA-dependent adaptations that promote DOPr expression on CINs all abolish PIT (7, 15). Together, these findings support a model in which BLA-mediated Pavlovian learning engages the BLA*→*NAc-S pathway to redundantly encode those associations in the enduring DOPr-dependent plasticity localized to NAc-S CINs. This plasticity appears, therefore, to establish a form of specific stimulus-outcome memory that is not required for Pavlovian learning or conditioned responding per se but is necessary for predictive stimuli subsequently to influence choice between instrumental actions during PIT.

Nevertheless, despite this extensive evidence for the involvement of the BLA*→*NAc-S pathway in encoding the specific S-O associations necessary for PIT, the circuit engaged during the retrieval of those associations remains unknown. Many contemporary theories of memory propose that memory encoding and retrieval depend on the same circuit; indeed, on the same neurons within circuits localized to a specialized memory store: so-called engram neurons (18). However, other views, most notably proceduralist theories of memory (19), are less prescriptive, arguing that event memories are localized not to a specialized store or cell but to the circuits and cell types engaged during their encoding and retrieval, which may consequently differ. The aim of the current series of experiments was, therefore, to determine whether the circuit that encodes the S-O associations necessary for PIT is also necessary for their retrieval; that is, whether the BLA*→*NAc-S circuit and the associated cell type-specific changes induced during S-O encoding also mediate the retrieval of those associations for specific PIT.

## RESULTS

### BLA-dependent changes in the NAc-S are sufficient to retrieve specific S-O associations for PIT

If the encoding and retrieval of specific S-O associations for PIT depends on the same circuit, then the changes induced in NAc-S CINs during encoding should be reinstated when those associations are retrieved, and that reinstatement should be necessary for PIT. In our first series of experiments, we sought, therefore, to assess whether the changes in the NAc-S induced during initial S-O exposure are recapitulated when subjects are re-exposed to those associations at retrieval, whether that recapitulation is sufficient to induce PIT and whether this process depends on the BLA. To address these questions, we capitalized on previous research showing that the effect of S-O exposure on DOPr-expression in NAc-S CINs can be disrupted by the DOPr agonist SNC80 (7). SNC80 causes DOPr internalization for 24 to 48 h and blocks the influence of predictive learning on choice, an effect that recovers as DOPr return to the membrane. Although this trafficking of internalized DOPr is often characterized as a spontaneous consequence of de-phosphorylation (20, 21), we hypothesized that, at least in the NAc-S, it reflects the retrieval of prior predictive learning.

Given this suggestion, we first investigated whether, after injection with SNC80, re-exposure to the trained S-O contingencies was sufficient to drive DOPrs back to the membrane, and, if so, on what timescale (Figure 1A). DOPr-eGFP mice underwent Pavlovian conditioning in which two stimuli (S1 and S2; white noise or tone) predicted two distinct food outcomes (O1 and O2; grain pellets or sucrose solution). At the end of the final training day, mice received a systemic injection of either SNC80 (snc), to internalize accumulated membrane DOPrs, or vehicle (veh). Seventeen hours later, half of the animals in each pharmacological condition were given a single Pavlovian reminder session (rem), while the others were handled (no). Brains were taken immediately afterwards for DOPr quantification, that is, before the usual recovery window, allowing us to isolate any experience-driven contribution to receptor translocation. During Pavlovian training (Figure 1B), mice successfully learned that the stimuli predicted the two outcomes: the number of magazine entries during stimulus presentations was higher than during their absence, as indicated by an elevation ratio significantly greater than 0.5. Importantly, we found that DOPr expression was greater in both vehicle groups and in SNC80-treated mice that received a reminder than in SNC-80 treated mice that did not (Figures 1C-D), confirming that a single reexposure to outcome-specific predictive learning is sufficient to reinstate DOPrs on NAc-S CINs. Furthermore, because this effect was observed immediately after the reminder and at a time well before spontaneous return, it identifies predictive learning itself as the active driver of DOPr trafficking to the somatic membrane.

**Fig. 1.**
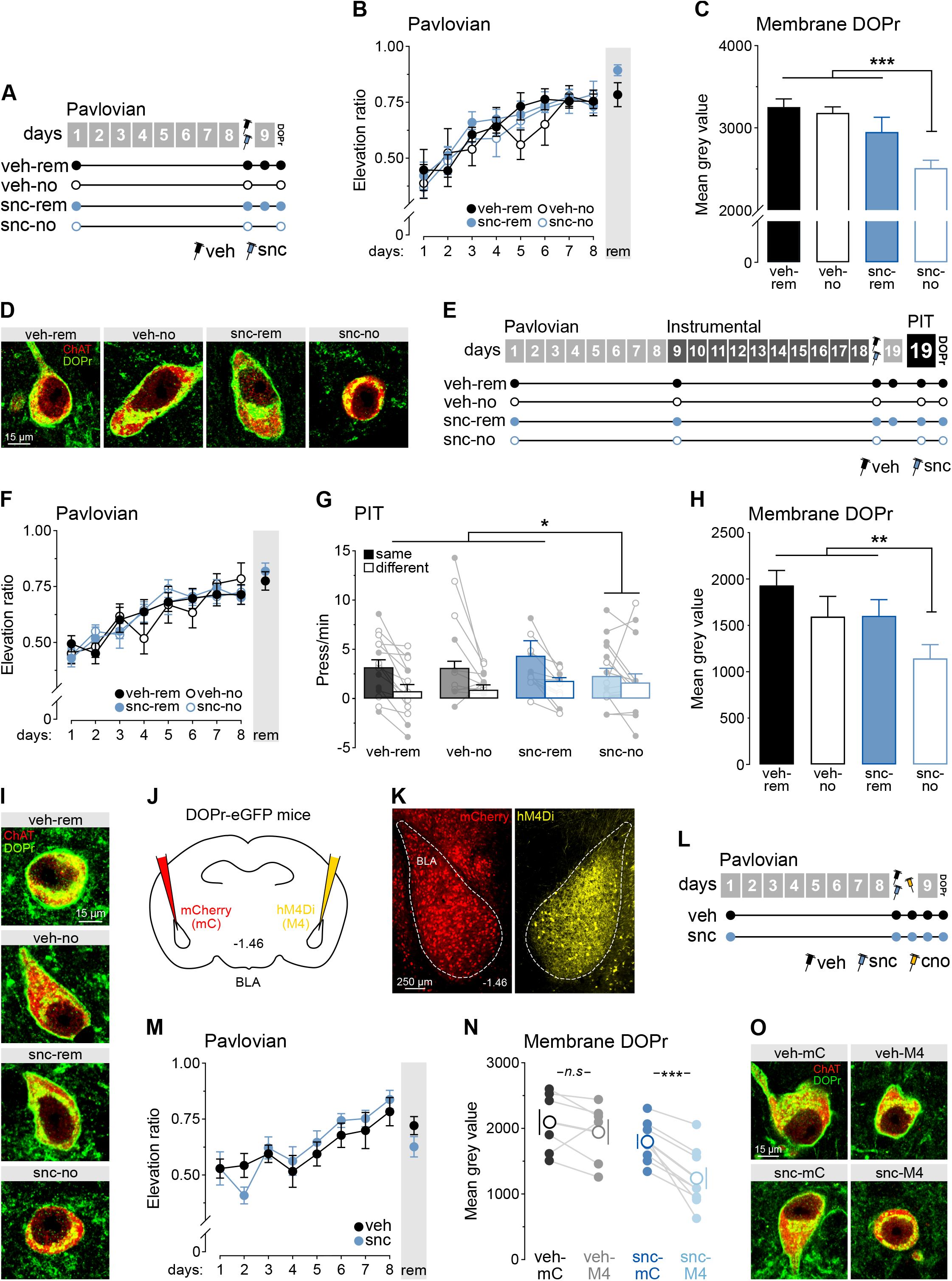
BLA-dependent changes in the NAc-S are sufficient to retrieve specific S-O associations for PIT. **(A)** Four groups of DOPr-eGFP mice were used: veh-rem (*n*=7; 3 M, 4 F); veh-no (*n*=6; 3 M, 3 F); snc-rem (*n*=7; 3 M, 4 F) and snc-no (*n*=7; 3 M, 4 F). Mice received Pavlovian training followed by SNC80 or vehicle injection. Half of the animals in each pharmacological condition were then given a retraining session, and DOPr expression levels were quantified. **(B)** Pavlovian training generated elevation ratios significantly above 0.5. Ratios were similar across groups (Fs<0.11) and increased over training (F(1,23)=89.34; p<0.001), with no group differences (Fs<0.04). Baseline responding did not differ (Fs<2.8). **(C)** Both vehicle-treated groups exhibited high levels of DOPr expression on NAc-S CINs (veh-rem vs. veh-no; F<0.17). These levels were similar in SNC80-treated mice that received additional S-O training (veh-rem + veh-no vs. snc-rem: F<3.21). DOPr expression was significantly lower in SNC80-treated mice that did not receive retraining only (veh-rem + veh-no + snc-rem vs. snc-no: F(1,23)=19.70; p<0.001). **(D)** Representative micrographs of membrane DOPr distribution on NAc-S CINs. **(E)** Four groups of DOPr-eGFP mice were used: veh-rem (*n*=15; 7 M, 8 F); veh-no (*n*=11; 6 M, 5 F); snc-rem (*n*=12; 7 M, 5 F) and snc-no (*n*=16; 10 M, 6 F). Mice received Pavlovian and instrumental training followed by SNC80 or vehicle injection. Half of the animals in each group were then given a Pavlovian retraining session. Next, all mice completed an outcome-specific PIT test followed by DOPr quantification. **(F)** Pavlovian training generated elevation ratios significantly above 0.5. Ratios were similar across groups (Fs<0.002) and gradually increased over sessions (F(1,50)=131.60; p<0.001) irrespective of group (Fs<0.01). Baseline responding was equivalent in all groups (Fs<1.3). **(G)** At test, both vehicle-treated groups (veh-rem: F(1,14)=23.84; p<0.001; veh-no: F(1,10)=5.04; p<0.05; veh-rem vs. veh-no: F<1.11) and the SNC80 group given S-O reminder (rem) displayed the outcome-specific PIT effect (snc-rem: F(1,11)=21.86; p<0.001; veh-rem + veh-no vs. snc-rem: F<0.3). This effect was absent in the other SNC80 group for which retraining of the S-O associations was omitted (snc-no: F<0.79; veh-rem + veh-no + snc-rem vs. snc-no: F(1,50)=4.13; p<0.05). Baseline responding (Fs<0.004) and overall lever press rates were similar between groups (Fs<0.23). Individual data points for females (open circles) and males (filled circles) are presented. **(H)** Both vehicle-treated groups exhibited high levels of DOPr expression on NAc-S CINs (veh-rem vs. veh-no: F<1.78). These levels were similar to those in SNC80-treated mice that received additional S-O training (veh-rem + veh-no vs. snc-rem: F<0.53). DOPr expression was significantly lower in SNC80-treated mice that did not receive retraining (veh-rem + veh-no + snc-rem vs. snc-no: F(1,50)=8.97; p<0.01). **(I)** Representative micrographs of membrane DOPr distribution on NAc-S CINs. **(J)** DOPr-eGFP mice received infusions of mCherry in one BLA and hM4Di in the contralateral BLA. **(K)** Representative micrographs showing viral spread in the BLA. **(L)** DOPr-eGFP mice were assigned to two pharmacological groups (veh or snc), combined with within-animal hemisphere counterbalancing of treatments across hemispheres. This yielded four conditions: Group veh (*n*=6; 3 M, 3 F): veh-mC and veh-M4; and Group snc (*n*=8; 6 M, 2 F): snc-mC and snc-M4. After recovery, all mice received Pavlovian training followed by SNC80 or vehicle. The next day, they received CNO and were retrained before assessing membrane DOPr levels. **(M)** Pavlovian training produced similar elevation ratios in both groups (F<0.25). Ratios increased over training (F(1,12)=36.28; p<0.001), with no group differences (F<1.45). Baseline responding was equivalent (F<0.3). **(N)** Overall DOPr expression on NAc-S CINs was higher in vehicle-than SNC80-treated mice (F(1,12)=5.32; p<0.05) and differed between the hemisphere infused with hM4Di vs. mCherry (F(1,12)=16.51; p<0.001), producing a treatment x hemisphere interaction (F(1,12)=5.38; p<0.05). DOPr levels in the hemisphere in which the BLA was treated with mCherry were significantly lower than in the hemisphere in which the BLA was treated with hM4Di (snc-mC vs. snc-M4: F(1,7)=25.49; p<0.001). **(O)** Representative micrographs of membrane DOPr distribution on NAc-S CINs. Data are shown as mean ± SEM. Asterisks denote significant effect (*: p<0.05; **: p<0.01; ***: p<0.001; *n*.*s*.: non-significant).

We next asked whether this retrieval-induced membrane DOPr expression also rescues the outcome-specific PIT effect typically abolished by SNC80 (Figure 1E). DOPr-eGFP mice underwent Pavlovian and instrumental conditioning, then received an SNC80 (snc) or vehicle (veh) injection. Seventeen hours later, half of the mice in each pharmacological condition received a single Pavlovian reminder session (rem) whereas the reminder were simply handled (no). Two hours later, the influence of predictive learning on choice was assessed in all mice in a PIT test, followed immediately by DOPr quantification. During Pavlovian training (Figure 1F), mice successfully learned that the two stimuli predicted the two outcomes. Instrumental training proceeded as expected (Figure S1A) and mice learned to press one lever to earn one outcome and to press another lever to obtain the other outcome. At test (Figure 1G), the vehicle-treated mice and SNC80-treated mice that received a reminder showed successful outcome-specific PIT effect, with the stimuli biasing performance toward the lever press action earning the same outcome as they predicted, relative to the action delivering the different outcome. By contrast, SNC80-treated mice that did not receive a reminder failed to show this effect. Furthermore, this failure was accompanied by low levels of membrane DOPr expression (Figures 1H-I), whereas the other three groups exhibited comparably higher DOPr expression. These results demonstrate that retrieval of Pavlovian S-O associations is sufficient to rescue outcome-specific PIT after SNC80 administration and provides clear evidence that both the encoding and retrieval of predictive learning increase membrane DOPr accumulation and promotes PIT.

Finally, we tested whether the effect of retrieval on DOPr membrane expression after SNC80 depends on the BLA (Figures 1J-L). We used a within-subject design in which the BLA was inactivated in one hemisphere during the reminder session, allowing us to compare DOPr accumulation on NAc-S CINs in that hemisphere with accumulation in the untreated hemisphere. DOPr-eGFP mice received unilateral BLA infusions of inhibitory hM4Di (M4) in one hemisphere and control mCherry (mC) in the other (with side counterbalanced; Figure S1B). Mice then underwent Pavlovian training followed by SNC80 (snc) or vehicle (veh) treatment. Sixteen hours later, all mice were injected with CNO to produce unilateral inactivation of the BLA and 45 min after that, they received a single Pavlovian reminder (rem) session followed immediately by DOPr quantification. Initial Pavlovian training proceeded as expected (Figure 1M). In vehicle-treated animals, DOPr expression on NAc-S CINs (Figures 1N-O) was similar between the inactivated and intact hemispheres. In SNC80-treated mice, by contrast, the hemisphere in which the BLA was inhibited during the reminder session showed lower DOPr expression on NAc-S CINs than the contralateral intact hemisphere. Thus, BLA inactivation prevented retrieval-induced DOPr translocation on NAc-S CINs, indicating that this process requires BLA activity. Together with the preceding experiments, these findings are consistent with the view that both the encoding and retrieval of specific S-O associations for PIT engages the BLA*→*NAc-S pathway and the encoding-related changes in DOPr expression on CINs within the NAc-S.

### The BLA input to the NAc-S encodes but does not retrieve stimulus-specific predictions

Our initial findings suggest the encoding and retrieval of S-O associations in the NAc-S depend on the BLA*→*NAc-S pathway. Because we did not use a pathway-specific approach, however, they are also consistent with other interpretations.

Our next series of experiments asked, therefore, whether the BLA’s contribution to retrieving S-O associations is mediated specifically by its projection to the NAc-S. We first examined whether BLA activity is necessary to retrieve S-O associations during the PIT test (Figures 2A-D). We then adopted a pathway specific optogenetic approach to assess the role of the BLA*→*NAc-S pathway, both in encoding S-O associations during predictive learning (Figures 2G-L) and in retrieving them during the PIT test (Figures 2M-P).

**Fig. 2.**
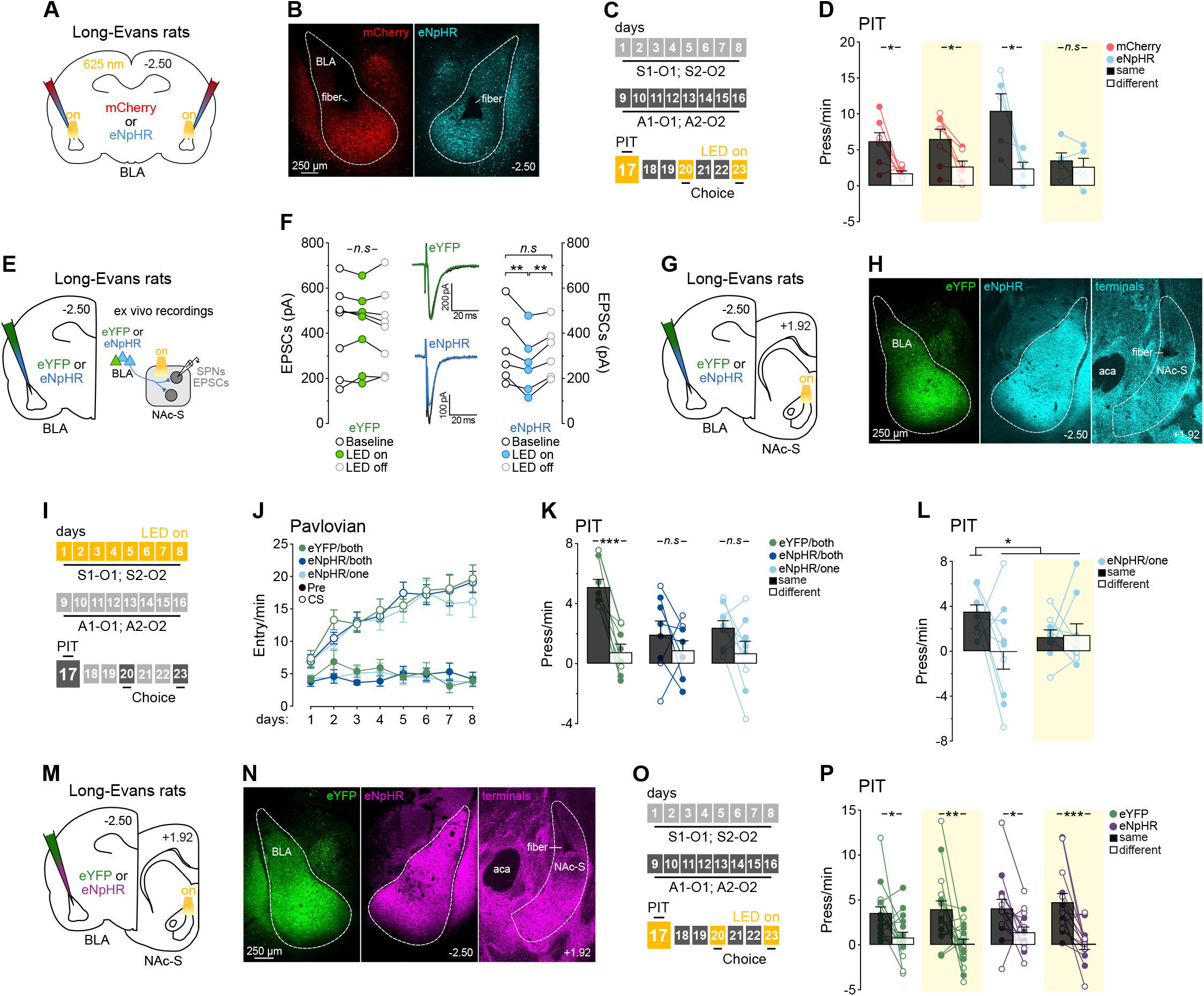
The BLA input to the NAc-S encodes but does not retrieve stimulus-specific predictions. **(A)** Long-Evans rats received bilateral mCherry or eNpHR infusions in the BLA and were implanted with fiber optics above the injection sites. **(B)** Representative micrographs showing the viral spread and fiber optic placement in the BLA. **(C)** Two groups of Long-Evans rats were used: mCherry (*n*=7; 4 M, 3 F) and eNpHR (*n*=5; 3 M, 2 F). After Pavlovian and instrumental training, they completed an outcome-specific PIT test in which LED illumination (625 nm, continuous) was delivered during half of the stimulus presentations. The following days, rats received two valued-based choice tests in which LED light was delivered during half of the trials (625 nm, continuous). **(D)** Silencing the BLA eliminated outcome-specific PIT. Baseline responding (F<1.01) and overall lever press rates during the test were similar between groups (F<0.32) and light conditions (F<1.93), with no group x light interaction (F<4.13). Lever press rates were higher on the action earning the same outcome as the stimuli compared to the action earning the different outcome (F(1,10)=20.21; p<0.001), regardless of group (F<0.03). There was a light x lever interaction (F(1,10)=9.07; p<0.01), and critically, a light x lever x group interaction during the presentation of the predictive stimuli (F(1,10)=6.45; p<0.05). Follow-up analyses revealed a successful transfer effect for the mCherry group whether the LED light was off (F(1,6)=9.43; p<0.05) or on (F(1,6)=8.07; p<0.05). By contrast, rats in the eNpHR group displayed outcome-specific PIT when the light was off (F(1,4)=10.37; p<0.05) but not on (F<0.73). Individual data points for females (open circles) and males (filled circles) are presented. Yellow rectangles represent LED light delivery at test. **(E)** Long-Evans rats received bilateral eYFP or eNpHR infusions in the BLA and recordings were made ex vivo from NAc-S spiny projection neurons. During electrical stimulation of BLA terminals, we applied 625 nm LED light to inhibit glutamate release (see methods). Evoked EPSCs were measured from NAc-S SPNs before (Baseline), during (LED on) and after (LED off) light delivery. This generated two groups: eYFP (*n*=3; 3 F) and eNpHR (*n*=4; 4 F). **(F)** In the eYFP group (left), LED light presentation had no effect on EPSCs amplitude (Fs<0.01). LED light delivered onto eNpHR-expressing BLA terminals (right) significantly reduced EPSCs amplitude in NAc-S SPNs, consistent with the inhibition of glutamate release (Baseline vs. LED on: F(1,5)=15.62, p<0.01; LED on vs. LED off: F(1,5)=26.32, p<0.01). EPSCs amplitude did not differ before and after light delivery (Baseline vs. LED off: F<0.5). Representative evoked EPSCs traces before (black) and during (green or blue) LED delivery are shown for an eYFP control (middle top) and an eNpHR rat (middle bottom). **(G)** Long-Evans rats received bilateral eYFP or eNpHR infusions in the BLA and were implanted with 1161 fiber optics above the NAc-S. **(H)** Representative micrographs showing the viral spread in the BLA (left and middle) and fiber optic placement in the NAc-S (right). **(I)** Three groups of Long-Evans rats were used: eYFP/both (*n*=8; 5 M, 3 F), eNpHR/both (*n*=8; 5 M, 3 F), and eNpHR/one (*n*=9; 6 M, 3 F). Rats in the eYFP and the eNpHR/both groups underwent Pavlovian training in which LED light delivery (625 nm, continuous) was paired with every S-O presentation. Rats in the reminder eNpHR/one group underwent Pavlovian training in which LED light delivery was paired with only one of the S-O presentation. All rats then received instrumental training, a PIT test, and two valued-based choice tests. **(J)** During Pavlovian conditioning, magazine entry rates were similar across groups (Fs<0.47) and increased over training (F(1,22)=41.66; p<0.001), with no group differences (Fs<1.28). Rates were higher in the presence of the stimuli than in their absence (F(1,22)=359.24; p<0.001), irrespective of group (Fs<0.78). The discrimination between the two periods increased as training progressed (F(1,22)=136.53; p<0.001), regardless of group (Fs<1.23). (K) Silencing the BLA*→*NAc-S pathway during predictive learning eliminated outcome-specific PIT. Baseline responding (Fs<1.13) during the test were similar between groups. Overall lever press rates were greater in the eYFP/both group relative to the eNpHR groups (eYFP/both vs. eNpHR/both + eNpHR/one: F(1,22)=5.60; p<0.05), with no difference between eNpHR groups (eNpHR/both vs. eNpHR/one: F<0.03). Rates were higher on the action earning the same outcome as the stimuli compared to the action earning the different outcome (F(1,22)=20.88; p<0.001), with a group x lever interaction during the presentation of the predictive stimuli (eYFP/both vs. eNpHR/both + eNpHR/one: F(1,22)=7.20; p<0.05). No group x lever interaction was detected between eNpHR groups (eNpHR/both vs. eNpHR/one: F<0.28). Follow-up analyses revealed a successful transfer effect for the eYFP/both group (F(1,7)=40.30; p<0.001), but not for the eNpHR/both group (F<0.96) nor the eNpHR/one group (F<3.68). Individual data points for females (open circles) and males (filled circles) are presented. **(L)** A separate set of contrasts was used to analyse performance for eNpHR/one group alone. A first within-subject contrast revealed no difference between “same” and “different” responding when the presented stimuli at test were paired with LED light delivery during Pavlovian training (same on vs. different on: F<0.02). A second within-subject contrast revealed no difference between these two conditions and “different” responding when the presented stimuli at test were paired with no LED light delivery during Pavlovian training (same on + different on vs. different off: F<0.84). A final contrast revealed an interaction between these three conditions and “same” responding when the presented stimuli at test were paired with the light off during Pavlovian training (same on + different on + different off vs. same off: (F(1,8)=6.59; p<0.05). Individual data points for females (open circles) and males (filled circles) are presented. The yellow rectangle represents LED light delivery that was paired with one of the S-O association during Pavlovian conditioning. **(M)** Long-Evans rats received bilateral eYFP or eNpHR infusions in the BLA and were implanted with optic fibers above the NAc-S. **(N)** Representative micrographs showing the viral spread in the BLA (left and middle) and fiber optic placement in the NAc-S (right). **(O)** Two groups of Long-Evans rats were used: eYFP (*n*=17; 10 M, 7 F) and eNpHR (*n*=14; 7 M, 7 F). After Pavlovian and instrumental training, they completed an outcome-specific PIT test in which LED illumination (625 nm, continuous) was delivered during half of the stimulus presentations. The following days, rats received two valued-based choice tests in which LED light (625 nm, continuous) was delivered during half of the trials. **(P)** Silencing the BLA*→*NAc-S pathway left outcome-specific PIT intact. Baseline responding (F<0.41) and overall lever press rates during the test were similar between groups (F<0.54) and light conditions (F<0.55), with no group x light interaction (F<0.09). Lever press rates were higher on the action earning the same outcome as the stimuli compared to the action earning the different outcome (F(1,29)=31.48; p<0.001), regardless of group (F<0.11). There was no light x lever interaction (F<3.01), and no light x lever x group interaction during the presentation of the predictive stimuli (F<0.27). Simple effect analyses revealed that lever press rates were higher on the same lever compared to the different lever in both groups whether the LED was off or on (eYFP off: F(1,16)=7.23; 1214 p<0.05; eYFP on: F(1,16)=9.85; p<0.01; eNpHR off: F(1,13)=6.64; p<0.05; eNpHR on: F(1,13)=19.78; p<0.001). Individual data points for females (open circles) and males (filled circles) are presented. Yellow rectangles represent LED light delivery at test. Data are shown as mean ± SEM. Asterisks denote significant effect (*: p<0.05; **: p<0.01; ***: p<0.001; *n*.*s*.: non-significant).

To examine whether BLA activity is required for choice at the time of the PIT test, we used optogenetics to inhibit that activity during stimulus presentations (Figures 2A-C). To target BLA glutamatergic projection neurons, rats received bilateral infusions of CaMKIIa-mCherry (mCherry) or CaMKIIa-eNpHR (eNpHR) in the BLA, with optic fibers implanted above the injection sites (Figure S2A). After Pavlovian (Figure S2B) and instrumental training (Figure S2C), rats completed a PIT test in which LED illumination (625 nm, continuous) was delivered during half of the stimulus presentations, counterbalanced across stimuli and order, enabling a within-subjects comparison of light-off and light-on trials. At test (Figure 2D), mCherry rats exhibited intact PIT across all trials. In contrast, eNpHR rats showed PIT only on light-off trials. On light-on trials, PIT was eliminated. To establish that these effects were specific to the influence of predictive learning on choice, all the rats were next given two outcome devaluation tests assessing their ability to use changes in outcome value to modify choice. The two value-based choice tests were conducted in which LED light was delivered during half of the test period (Figure S2D). LED light delivery had no effect on performance in the choice tests in either group, indicating that BLA silencing at test selectively disrupts the influence of predictive stimuli on choice between actions.

We next aimed to establish whether these effects of silencing the BLA depend on its projection to the NAc-S, by inhibiting BLA terminals in the NAc-S during either the encoding or the retrieval of predictive learning for PIT. We first validated the approach, confirming that the inhibitory opsin eNpHR provided reliable, terminal-specific inhibition of this pathway (Figures 2E-F) producing precise, reversible inhibition of the BLA*→*NAc-S pathway, suitable for our behavioral manipulations. Using this approach (Figures 2G-I) we first sought to replicate our previous observation that activity in the BLA*→*NAc-S pathway during Pavlovian conditioning is necessary to encode outcome-specific predictions that later influence choice between actions (7). Three groups of rats received bilateral BLA infusions – one with the control eYFP virus and two with eNpHR – and optic fibers were implanted above the NAc-S (Figure S2E). After recovery, all groups underwent Pavlovian training with LED illumination (625 nm, continuous) delivered during stimulus-outcome pairings. In one eNpHR group, illumination was given during only one of the pairings; either S1-O1 or S2-O2, counterbalanced for stimulus and outcome (eNpHR/one). For the eYFP and remaining eNpHR group, illumination was delivered during both pairings (eYFP/both: eNpHR/both). We predicted finding normal PIT in eYFP controls. In contrast, whereas the eNpHR/one group should show PIT during the stimulus trained with LED off but not that trained with LED on, the eNpHR/both group should fail to show PIT during both stimuli. To test these predictions, rats completed instrumental training (Figure S2F) followed by a single outcome-specific PIT test. Pavlovian training (Figure 2J) revealed that silencing the BLA*→*NAc-S pathway did not alter conditioned responding during acquisition of outcome-specific predictions. Rats then completed instrumental training (Figure S2F), followed by a single PIT test. Whereas eYFP control rats displayed PIT, with stimulus presentation biasing performance toward the action earning the same outcome (Figure 2K), inhibiting the BLA*→*NAc-S pathway during predictive learning selectively abolished this effect. In the eNpHR/both group, the bias was abolished for both stimuli. In the eNpHR/one group, it was abolished for the stimulus trained under illumination but preserved for the stimulus trained without it (Figure 2L). Finally, these effects were specific to the influence of predictive learning on choice: rats retained an intact capacity to use outcome value to choose between these two levers in an outcome devaluation test (Figure S2G).

We next tested whether the BLA*→*NAc-S pathway also supports the retrieval of specific S-O associations at test. If the BLA encodes and updates the current predictive validity of Pavlovian stimuli, it may also directly activate the neural infrastructure established in the NAc-S during predictive learning (i.e., the DOPr-based memory) to retrieve those associations and bias action selection. Accordingly, we silenced the BLA*→*NAc-S pathway during the PIT test, using the within-subjects approach described above (Figures 2M-O). Rats were infused bilaterally in the BLA with eYFP or eNpHR, with optic fibers implanted above the NAc-S (Figure S2H). All rats received Pavlovian conditioning (Figure S2I), with each of the two stimuli associated with a different outcome, followed by instrumental conditioning (Figure S2J) in which they were trained to press two levers for those same outcomes before they were given the PIT test (Figure 2P). LED illumination (625 nm, continuous) was delivered during half of the stimulus presentations, enabling a within-subjects comparison of light-off and light-on trials. Silencing had no effect on PIT: predictive stimuli biased choice between actions whether the BLA*→*NAc-S pathway was silenced or not (Figure S2K).

Together, these experiments reveal a striking dissociation: the BLA*→*NAc-S pathway is necessary when encoding the specific stimulus-outcome associations but is dispensable during retrieval. Silencing the pathway during one S-O association affected the encoding of that association without affecting encoding of the other (Figure 2L). Yet the same pathway was not required for retrieval of predictive learning at the time of choice (Figure 2P), even though BLA activity itself is required (Figure 2D). This suggests that the BLA must control choice at test via an alternative pathway.

### A novel amygdala-cortico-striatal circuit retrieves outcome-specific predictions for choice

The infralimbic cortex (IL) maintains reciprocal connections with the BLA (22–24) and projects to the NAc-S (25, 26). It therefore emerges as a candidate intermediary, capable of relaying BLA-mediated predictive learning to control NAc-S output for choice. Consistent with this, disconnecting the IL from the NAc-S using asymmetrical lesions abolishes PIT (27). We therefore hypothesized that the IL mediates the influence of the BLA on choice at test. To evaluate this circuit, we first conducted tracing studies to establish whether BLA neurons projecting to the NAc-S also send collaterals to the IL, or whether the two form independent circuits (Figures 3A-B). Rats received unilateral retrograde eGFP or mCherry injections into the IL and ipsilateral NAc-S, labelling BLA neurons projecting to each target and identifying dual-projecting cells. Spatial distributions (Figure 3C) across the rostrocaudal BLA revealed greater posterior contributions to both regions. Of all labelled BLA neurons, more projected to the NAc-S (*≈* 64 %) than the IL (*≈* 33 %), and few were double-labelled (*≈* 3 %), indicating largely distinct projection populations (Figure 3D). Labelling densities did not differ between tracers for the IL (Figure 3E) or the NAc-S (Figure 3F), indicating comparable infection efficiency. The BLA therefore maintains largely separate projections to the NAc-S and IL, consistent with the possibility that these projections serve distinct functions in PIT.

**Fig. 3.**
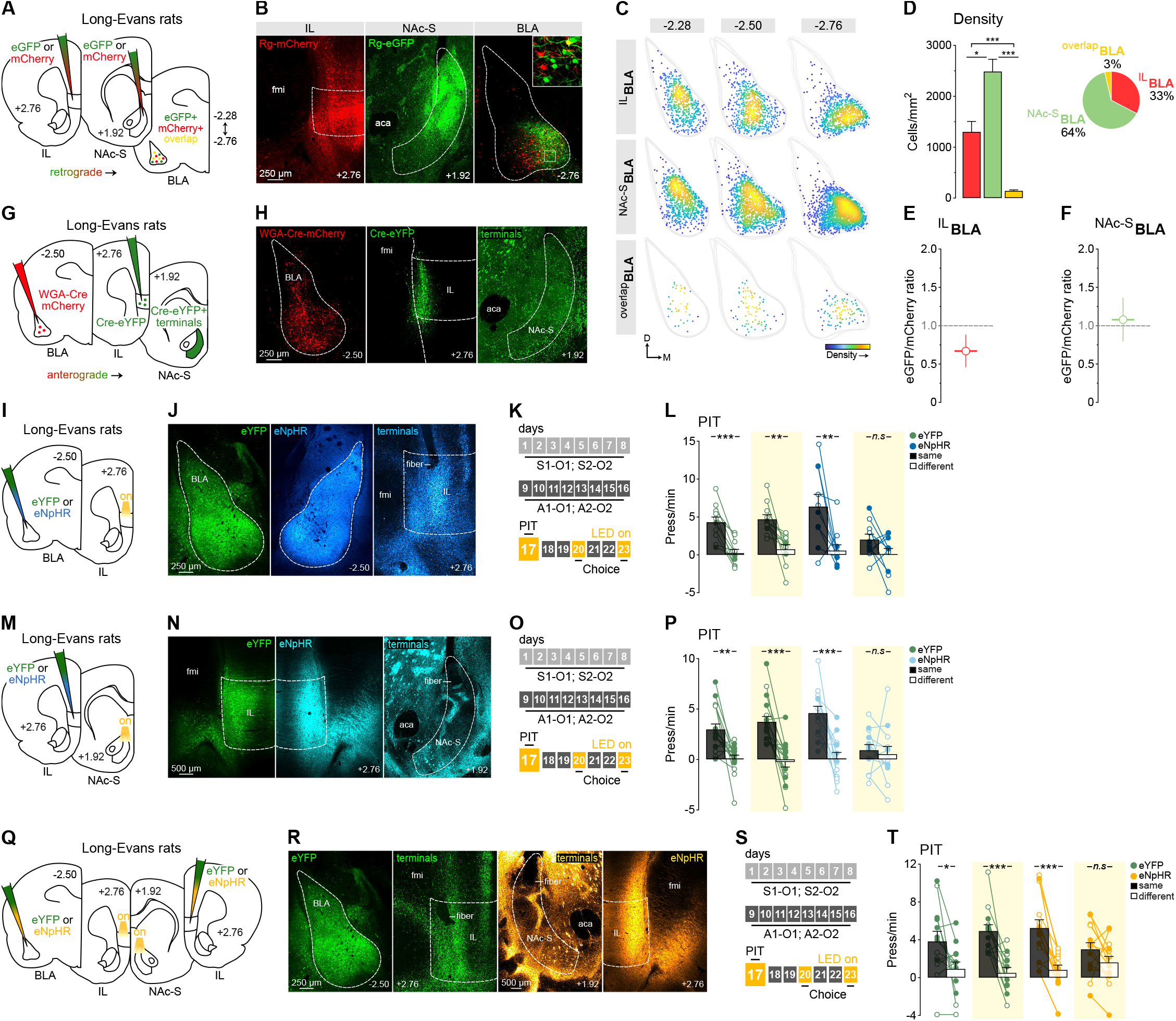
A novel amygdala-cortico-striatal circuit retrieves outcome-specific predictions for choice. **(A)** Long-Evans rats received unilateral retrograde eGFP or mCherry in the IL and ipsilateral NAc-S (*n*=7; 7 M; counterbalanced viruses and hemispheres). **(B)** Representative micrographs showing the viral spread in the IL (left), ipsilateral NAc-S (middle), and infected cell bodies in the BLA (right). Inset shows zoomed BLA-infected cell bodies with eGFP (green), mCherry (red), or both viruses (yellow; top right). **(C)** Scatterplots showing separate spatial distributions of BLA-infected cell bodies at 3 different bregma coordinates, with pseudocolored densities highlighting clustering (stacked hemisections). 28 hemisections in 7 rats were used (9-10 sections per bregma; 1-2 sections per animal). **(D)** Density of BLA neurons projecting to the IL (red), NAc-S (green) or both regions (yellow), averaged across 3 bregma levels (left). BLA projections to the NAc-S predominated (*≈* 64 %; right) and were significantly denser than the projections to the IL (*≈* 33 %; Dunn: Z=2.52; p<0.05) and to the overlapping population (*≈* 3 %; Dunn: Z=7.34; p<0.001). Projections to the IL were also significantly denser than the overlap (Dunn: Z=4.82; p<0.001), indicating minimal convergence between projection populations. **(E)** Injections of eGFP and mCherry viruses into the IL resulted in comparable patterns and densities of labelled BLA neurons, with a mean eGFP/mCherry density ratio of *≈* 0.67. No difference was observed between tracers (Mann-Whitney: U=68; p=0.24). **(F)** Injections in the NAc-S produced a mean eGFP/mCherry density ratio close to unity *≈* 1.08, with no detectable difference in mean labelling density between eGFP- and mCherry-injected animals (Welch: t=0.28; p=0.78). **(G)** Long-Evans rats received unilateral WGA-Cre-mCherry infusions in the BLA and ipsilateral Cre-eYFP infusions in the IL (*n*=4; 4 F; hemispheres counterbalanced). **(H)** Representative micrographs showing the viral spread in the BLA (left), eYFP expression in postsynaptic IL neurons that had accumulated WGA-Cre (middle), and the presence of eYFP-positive terminals in the NAc-S (right). (I) Long-Evans rats received bilateral eYFP or eNpHR infusions in the BLA and were implanted with fiber optics above the IL. **(J)** Representative micrographs showing the viral spread in the BLA (left and middle) and fiber optic placement in the IL (right). **(K)** Two groups of Long-Evans rats were used: eYFP (*n*=11; 2 M, 9 F) and eNpHR (*n*=10; 4 M, 6 F). After Pavlovian and instrumental training, they completed an outcome-specific PIT test in which LED illumination (625 nm, continuous) was delivered during half of the stimulus presentations. The following days, rats received two value-based choice tests in which the LED light was delivered during half of the trials (625 nm, continuous). **(L)** Silencing the BLA*→*IL pathway impaired outcome-specific PIT. Baseline responding (F<2.31) and overall lever press 1256 rates during the test were similar between groups (F<0.18) and light conditions (F<3.54). However, a group x light interaction emerged (F(1,19)=7.42; p<0.05). Lever press rates were higher on the action earning the same outcome as the stimuli compared to the action earning the different outcome (F(1,19)=28.43; p<0.001), regardless of group (F<0.01). There was a light x lever interaction (F(1,19)=5.21; p<0.05), and critically, a light x lever x group interaction during the presentation of the predictive stimuli (F(1,19)=4.57; p<0.05). Follow-up analyses revealed a successful transfer effect for the eYFP group whether the LED light was off (F(1,10)=38.92; p<0.001) or on (F(1,10)=15.67; p<0.01). By contrast, rats in the eNpHR group displayed outcome-specific PIT when the light was off (F(1,9)=9.80; p<0.01) but not on (F<2.94). Individual data points for female (open circles) and male (filled circles) are presented. Yellow rectangles represent LED light delivery at test. **(M)** Long-Evans rats received bilateral eYFP or eNpHR infusions in the IL and were implanted with fiber optics above the NAc-S. **(N)** Representative micrographs showing the viral spread in the IL (left and middle) and fiber optic placement in the NAc-S (right). **(O)** Two groups of Long-Evans rats were used: eYFP (*n*=17; 9 M, 8 F) and eNpHR (*n*=13; 6 M, 7 F). After Pavlovian and instrumental training, they completed an outcome-specific PIT test in which LED illumination (625 nm, continuous) was delivered during half of the stimulus presentations. The following days, a subset of animals in each group (eYFP: *n*=10; 5 M, 5 F; eNpHR: *n*=8; 4 M, 4 F) received two value-based choice tests in which half of the trials were under LED light delivery (625 nm, continuous). **(P)** Silencing the IL*→*NAc-S pathway abolished outcome-specific PIT. Baseline responding (F<3.68) and overall lever press rates during the test were similar between groups (F<0.07) and light conditions (F<3.98). However, a group x light interaction emerged (F(1,28)=6.69; p<0.05). Lever press rates were higher on the action earning the same outcome as the stimuli compared to the action earning the different outcome (F(1,28)=33.79; p<0.001), regardless of group (F<0.98). There was a light x lever interaction (F(1,28)=4.63; p<0.05), and a light x lever x group interaction during the presentation of the predictive stimuli (F(1,28)=13.64; p<0.001). Follow-up analyses revealed a successful transfer effect for the eYFP group whether the LED light was off (F(1,16)=14.22; p<0.01) or on (F(1,16)=27.45; p<0.001). By contrast, rats in the eNpHR group displayed outcome-specific PIT when the light was off (F(1,12)=18.92; p<0.001) but not on (F<0.16). Individual data points for female (open circles) and male (filled circles) are presented. Yellow rectangles represent LED light delivery at test. **(Q)** Long-Evans rats received unilateral injections of either eYFP or eNpHR in the BLA in one hemisphere and in the IL in the contralateral hemisphere. Fiber optics were implanted above the IL on the BLA-infused side, and above the NAc-S on the IL-infused side. **(R)** Representative micrographs showing the viral spread in the BLA (left, green) and IL (right, orange), as well as fiber optic placement in the IL (middle left, green) and the NAc-S (middle right, orange). **(S)** Two groups of Long-Evans rats were used: eYFP (*n*=13; 7 M, 6 F) and eNpHR (*n*=14; 7 M, 7 F). After Pavlovian and instrumental training, they completed an outcome-specific PIT test in which LED illumination (625 nm, continuous) was delivered during half of the stimulus presentations. The following days, rats received two value-based choice tests in which half of the trials were under LED light delivery (625 nm, continuous). **(T)** Disconnecting the BLA*→*IL and IL*→*NAc-S pathways impaired outcome-specific PIT. Baseline responding (F<4.11) and overall lever press rates during the test were similar between groups (F<0.03) and light conditions (F<0.17), with no group x light interaction (F<1.22). Lever press rates were higher on the action earning the same outcome as the stimuli compared to the action earning the different outcome (F(1,25)=34.29; p<0.001), regardless of group (F<0.46). There was no light x lever interaction (F<1.19), but critically, there was a light x lever x group interaction during the presentation of the predictive stimuli (F(1,25)=10.60; p<0.01). Follow-up analyses revealed a successful transfer effect for the eYFP group whether the LED light was off (F(1,12)=6.05; p<0.05) or on (F(1,12)=22.97; p<0.001). By contrast, rats in the eNpHR group displayed outcome-specific PIT when the light was off (F(1,13)=29.39; p<0.001) but not on (F<3.05). Individual data points for female (open circles) and male (filled circles) are presented. Yellow rectangles represent LED light delivery at test. Data are shown as mean ± SEM. Asterisks denote significant effect (*: p<0.05; **: p<0.01; ***:p<0.001; *n*.*s*.: non-significant).

Next, to determine whether IL neurons targeted by the BLA project to the NAc-S, we conducted an anterograde transsynaptic tracing experiment (Figures 3G-H). Rats received unilateral infusions of an AAV expressing WGA-Cre in the BLA, together with a Cre-dependent eYFP vector in the ipsilateral IL. This approach allowed WGA-Cre to transfer from BLA terminals to postsynaptic IL neurons (28), driving eYFP expression selectively in IL neurons receiving BLA inputs. As shown in Figure 3H, eYFP-positive somata were present in the IL, and labelled terminals appeared in the NAc-S. Consistent with previous work, labelling was dense in IL layer 5 (22–24), with additional labelling in layer 6, a pattern also observed in primates (29). eYFP-positive terminals were found in the ipsilateral medial NAc-S but also in the nucleus accumbens core (NAc-C). This anatomical overlap does not by itself establish which target is functionally relevant, a question we address directly below (Figures S3K-L). These findings do, however, suggest that the NAc-S receives projections from IL neurons that are themselves targets of BLA inputs, providing evidence for a BLA*→*IL*→*NAc-S circuit.

We next investigated the causal role of this circuit in the retrieval of predictive learning during the PIT test, beginning with the BLA*→*IL arm. In our first experiment (Figures 3I-K), two groups of rats received bilateral infusions of either the eYFP or eNpHR in the BLA and were implanted with optic fibers above the IL (Figure S3A). After Pavlovian (Figure S3B) and instrumental conditioning (Figure S3C), we assessed the role of the BLA*→*IL pathway in the expression of PIT using the within-subjects procedure described above (Figure 3L). Bilateral silencing of BLA terminals in the IL at test abolished the rats’ ability to use predictive learning to guide choice between actions on light-on but not light-off trials. In contrast, eYFP controls displayed PIT under both light conditions, as did an anatomical control group of eNpHR rats with off-target virus or fiber placements in one hemisphere (Figure S3D). LED illumination had no effect on choice after outcome devaluation in either group (Figure S3E).

Having shown that BLA*→*IL projections are required for PIT, we next tested the second arm of the proposed circuit, asking whether the IL*→*NAc-S projection is also recruited during its expression (Figures 3M-O). Two groups of rats received bilateral infusions of either eNpHR or eYFP in the IL and were implanted with optic fibers above the NAc-S (Figure S3F). After Pavlovian (Figure S3G) and instrumental conditioning (Figure S3H), rats were tested using the within-subject procedure. Silencing IL terminals in the NAc-S abolished PIT in the eNpHR group (Figure 3P), whereas PIT was intact in eYFP controls and in an anatomical control group with off-target virus or fiber placements in one hemisphere (Figure S3I). Silencing IL terminals in NAc-S did not affect outcome devaluation in either group (Figure S3J).

Finally, because our tracing revealed considerable IL*→*NAc-C labelling (Figure 3H), we asked whether the effects on choice were specific to the IL*→*NAc-S pathway. We repeated the experiment, silencing the IL*→*NAc-C pathway at test using the same procedure. This had no effect, establishing that the PIT deficit is specific to the NAc-S (Figures S3K-L). Together, these results suggest that the BLA enables retrieval of specific stimulus-outcome associations for PIT expression by modulating IL activity. The IL in turn recruits the plasticity induced during S-O encoding, biasing choice between actions at test.

To provide a direct test of this hypothesis and so establish whether concurrent activity in the BLA*→*IL and IL*→*NAc-S pathways is required for the expression of PIT, we implemented a disconnection procedure in which the BLA*→*IL pathway was silenced in one hemisphere and the IL*→*NAc-S pathway was silenced in the contralateral hemisphere (Figures 3Q-S). The validity of this approach rests on our earlier findings that unilateral silencing of either pathway alone did not disrupt PIT (Figures S3D, I), indicating that a single intact hemisphere is sufficient to support choice. Two groups of rats received unilateral infusions of either eNpHR or the eYFP into the BLA of one hemisphere and into the IL of the contralateral hemisphere, with optic fibers implanted above the ipsilateral IL and NAc-S respectively, allowing terminal specific inhibition of each pathway (Figure S3M). Pavlovian (Figure S3N) and instrumental conditioning (Figure S3O) proceeded as expected. At test (Figure 3T), eYFP controls displayed intact PIT, whereas eNpHR rats did not. To confirm that these effects were specific to predictive learning, we conducted two value-based choice tests with matched light-on and light-off conditions (Figure S3P), finding that silencing did not affect the sensitivity of choice to outcome-devaluation. The results show that concurrent activity in both pathways is required to bias choice performance at test, supporting a model in which the BLA regulates the expression of specific S-O memories in the NAc-S by modulating IL activity at the time of choice.

### The BLA input to the IL mediates retrieval via NAc-S CINs

Our preceding experiments implicate a BLA*→*IL*→*NAc-S circuit in the expression of PIT at the time of choice. Whether this circuit, or parts of it, also operates during the encoding of predictive learning is less clear. We have found that the BLA*→*NAc-S circuit is required during encoding (Figures 2K-L) but not retrieval (Figure 2P), whereas the BLA*→*IL is required at retrieval (Figure 3L). However, whether this latter projection operates exclusively at retrieval remains unknown, nor is it known whether the IL*→*NAc-S circuit also influences DOPr expression on CINs. To address these questions, we first silenced BLA inputs to the IL selectively during Pavlovian training (Figures 4A-C). Rats were bilaterally infused with eYFP or eNpHR in the BLA and implanted with optic fibers above the IL (Figure S4A). After recovery, both groups underwent Pavlovian training in which LED illumination (625 nm, continuous) accompanied every S-O presentation. Rats subsequently acquired instrumental contingencies normally (Figure S4B) and completed a single PIT test followed by two value-based choice tests (Figure S4C). Conditioned responding during Pavlovian training was unaffected by silencing (Figure 4D). Critically, however, the manipulation impaired subsequent choice in the PIT test (Figure 4E) without affecting choice after outcome-devaluation (Figure S4C). BLA*→*IL signaling is therefore required during both the encoding and retrieval of S-O associations during PIT, unlike the BLA*→*NAc-S pathway, whose contribution is restricted to encoding (Figures 2K-L).

**Fig. 4.**
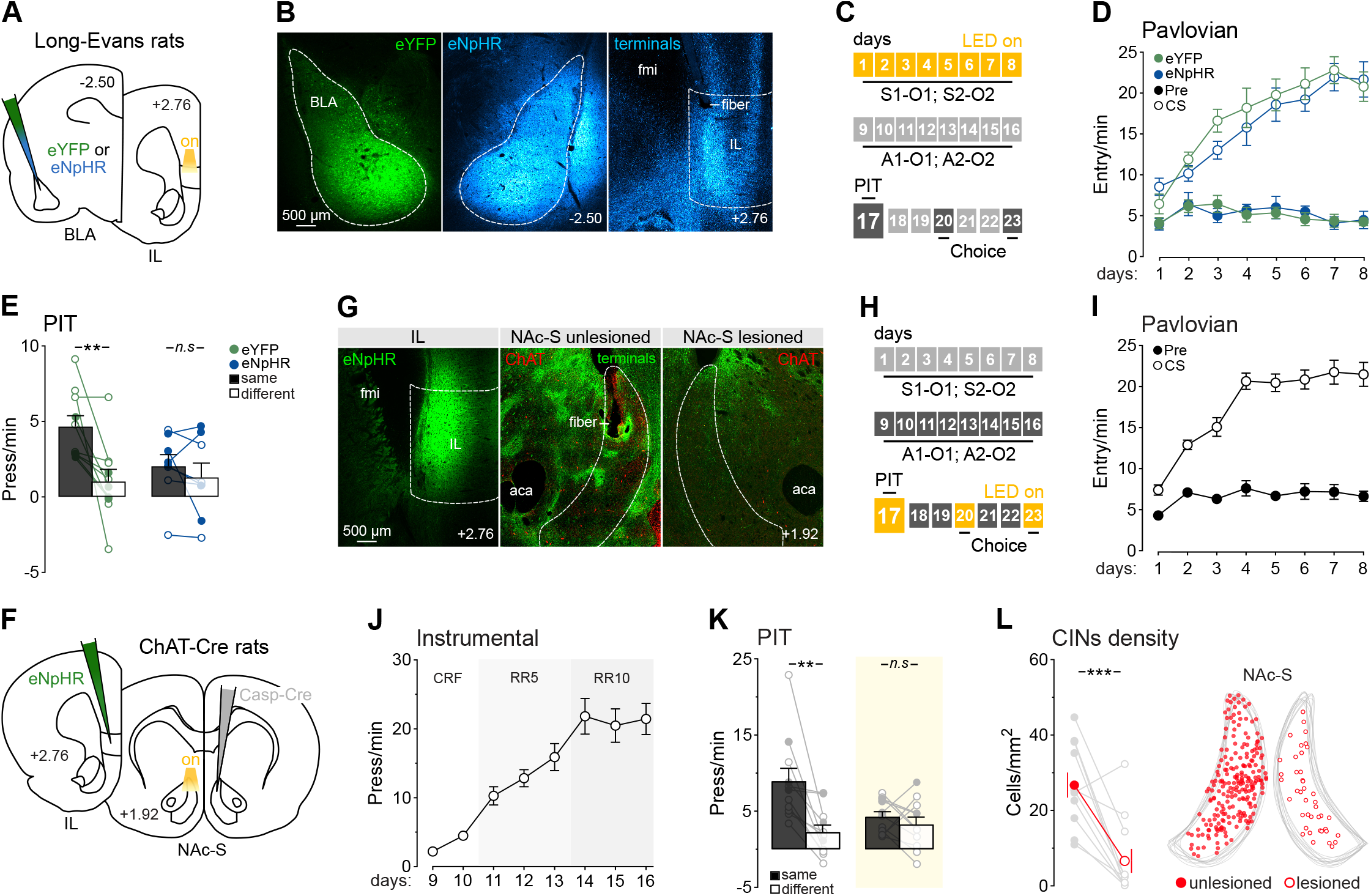
The BLA input to the IL mediates retrieval via NAc-S CINs. **(A)** Long-Evans rats received bilateral eYFP or eNpHR infusions in the BLA and were implanted with fiber optics above the IL. **(B)** Representative micrographs showing the viral spread in the BLA (left and middle) and fiber optic placement in the IL (right). **(C)** Two groups of Long-Evans rats were used: eYFP (*n*=10; 4 M, 6 F) and eNpHR (*n*=8; 4 M, 4 F). Rats underwent Pavlovian training in which LED light delivery (625 nm, continuous) was paired with every S-O presentation, followed by instrumental training, a PIT test, and two value-based choice tests. **(D)** During Pavlovian conditioning, magazine entry rates were similar across groups (F<0.22) and increased over training (F(1,16)=54.34; p<0.001), with no group differences (F<0.04). Rates were higher in the presence of the stimuli than in their absence (F(1,16)=249.64; p<0.001), irrespective of group (F<0.67). The discrimination between the two periods increased as training progressed (F(1,16)=158.14; p<0.001), regardless of group (F<0.03). **(E)** Silencing the BLA*→*IL pathway during predictive learning eliminated outcome-specific PIT. Baseline responding (F<0.05) and overall lever press rates during the test were similar between groups (F<1.51). Rates were higher on the action earning the same outcome as the stimuli compared to the action earning the different outcome (F(1,16)=11.41; p<0.01), with a group x lever interaction during the presentation of the predictive stimuli (F(1,16)=4.99; p<0.05). Follow-up analyses revealed a successful transfer effect for the eYFP group (F(1,9)=12.61; p<0.01), but not for the eNpHR group (F<1.23). Individual data points for females (open circles) and males (filled circles) are presented. **(F)** ChAT-Cre rats received unilateral eNpHR infusions in the IL and were implanted with a fiber optic above the ipsilateral NAc-S. They also received contralateral Caspase-Cre (Casp-Cre) infusions to specifically lesion NAc-S CINs (counterbalanced hemispheres). **(G)** Representative micrographs showing the viral spread in the IL (left), ipsilateral fiber optic placement in the NAc-S (middle), as well as ChAT staining in the unlesioned and lesioned NAc-S (middle and right). **(H)** One group of ChAT-Cre rats was used (*n*=11; 3 M, 8 F). After Pavlovian and instrumental training, they completed an outcome-specific PIT test in which LED illumination (625 nm, continuous) was delivered during half of the stimulus presentations. The following days, rats received two value-based choice tests in which half of the trials were under LED light delivery (625 nm, continuous). **(I)** Unilateral lesion of NAc-S CINs did not impact Pavlovian performance. Magazine entry rates increased over training (F(1,10)=72.87; p<0.001) and were higher in the presence of the stimuli than in their absence (F(1,10)=173.55; p<0.001). The discrimination between the two periods increased as training progressed (F(1,10)=48.64; p<0.001). **(J)** Unilateral lesion 1349 of NAc-S CINs did not impact instrumental performance. Lever press responding increased as training progressed (F(1,10)=77.52; p<0.001). **(K)** Unilateral silencing of the IL*→*NAc-S pathway combined with contralateral lesion of NAc-S CINs abolished outcome-specific PIT. Overall lever press rates during the test were similar between light conditions (F<4.70) and higher on the action earning the same outcome as the stimuli compared to the action earning the different outcome (F(1,10)=19.98; p<0.001). There was a light x lever interaction during the presentation of the predictive stimuli (F(1,10)=6.29; p<0.05). Follow-up analyses revealed a successful transfer effect when the light was off (F(1,10)=15.75; p<0.01) but not on (F<0.83). Individual data points for females (open circles) and males (filled circles) are presented. The yellow rectangle represents LED light delivery at test. **(L)** Average densities of CINs in the NAc-S between hemispheres that were intact (filled circles) and lesioned (open circles). Light gray dots are individual hemisections density averaged per animal. Red-coloured segments indicate the SEM (left). Caspase-Cre infusions successfully ablated NAc-S CINs (F(1,10)=21.71; p<0.001). Spatial distribution of CINs stained with ChAT in the NAc-S between hemispheres (stacked hemisections) that were intact (filled red circles) and lesioned (open red circles). 21 hemisections in 11 rats were used (1-2 sections per animal; right). Data are shown as mean ± SEM. Asterisks denote significant effects (**: p<0.01; ***: p<0.001; *n*.*s*.: non-significant).

Given these findings, we next evaluated the role of the IL in the changes induced in the NAc-S during Pavlovian conditioning; specifically, the accumulation of DOPr on CINs characterized above (Figures S4D-F). DOPr-eGFP mice received bilateral IL lesions or sham surgery before Pavlovian conditioning (Figure S4E), and DOPr expression on NAc-S CINs was compared between groups. IL lesion had no effect on DOPr expression (Figure S4F). Although these studies do not reveal what associative information the BLA*→*IL pathway carries during conditioning, they do indicate that it is not required for the plasticity conveyed to the NAc-S via the BLA*→*NAc-S pathway.

Nevertheless, these results suggest that the BLA facilitates the retrieval of predictive associations at test by re-engaging, via its projections to the IL, the cortical circuitry involved during encoding but leaves open how the retrieved information is read out by downstream structures to bias action selection. Based on the experiments above and our proposed model of PIT expression (4, 7, 13, 16, 17), we propose that the critical site of this readout is the NAc-S. In this model, cortical inputs to the NAc-S initiate neuromodulatory processes that activate DOPr, reducing acetylcholine release from CINs and thereby disinhibiting D1-spiny projection neurons (D1-SPNs) projecting to the ventral pallidum (30). Within this framework, IL projections to the NAc-S are well positioned to convey outcome-specific predictions retrieved from the BLA and, indirectly, to modulate CIN activity. Consistent with this, genetic ablation of NAc-S CINs abolishes PIT, with the magnitude of impairment scaling with the extent of CIN ablation (7).

To test this account and so establish whether both IL inputs to the NAc-S and local CIN signaling operate together to support PIT, we again used a within-subject functional disconnection procedure (Figures 4F-H). It exploited the fact that unilateral silencing of the IL*→*NAc-S pathway alone has no effect on PIT (Figure S3I) and that only complete bilateral CIN ablation abolishes PIT (7). By silencing the IL*→*NAc-S pathway in one hemisphere while ablating CINs in the contralateral NAc-S, we ensured that neither hemisphere retained sufficient information to induce PIT. To enable selective CIN ablation, this experiment used ChAT-Cre rats, in which Cre recombinase is expressed in CINs. Rats received unilateral eNpHR infusions in the IL with an optic fiber above the ipsilateral NAc-S, and a contralateral infusion of Caspase-Cre into the NAc-S to ablate local CINs (Figure 4L; Figure S4G). Following recovery, rats displayed normal Pavlovian (Figure 4I) and instrumental performance (Figure 4J). During the PIT test (Figure 4K), continuous LED illumination (625 nm) was delivered during half of the trials. On LED-off trials, PIT was intact, with the spared hemisphere (maintaining both an intact IL*→*NAc-S projection and intact CINs) supporting normal choice. In contrast, on LED-on trials, the now-complete disconnection abolished PIT, with rats distributing responding similarly across both actions during stimulus presentation. Finally, to confirm these effects were specific to the influence of predictive learning on choice, we conducted two value-based choice tests using matched LED conditions (Figure S4H) finding that choice after outcome-devaluation was unaffected by illumination.

## DISCUSSION

The finding that Pavlovian stimulus-outcome associations influence choice between instrumental actions has been important for associative learning theories, having sometimes been interpreted as evidence that Pavlovian and instrumental conditioning share a common representational structure. The current data provide evidence for a very different account: that Pavlovian and instrumental conditioning are essentially independent forms of learning whose interaction is mediated by a specialized circuit. We have characterized this circuit as producing a specialized form of memory, one driven by G protein-coupled receptor (GPCR) signaling that encodes stimulus-outcome associations independently of their encoding for Pavlovian conditioning and in a way that allows their retrieval to activate a broader circuit to influence instrumental performance. We have previously described the neural basis of this encoding process, demonstrating that BLA-induced plasticity at D1-SPNs in the NAc-S causes the release of substance P, which binds to neurokinin-1 receptors on CINs to induce the translocation of DOPr to the somatic membrane; a form of plasticity that reflects specific stimulus-outcome learning (7). However, to date, the neural processes mediating the retrieval of this memory during the transfer test have not been assessed.

The retrieval of specific S-O associations occurs many days after their encoding and so, for reasons of parsimony, we began by assuming that retrieval engages the same circuit and cell types engaged as those recruited during that initial process. We initially found some evidence concordant with that assumption, at least with respect to the role of the BLA in retrieving the changes in DOPrs on CINs at a time when those receptors were internalized. Contrary to that view, however, our subsequent experiments found that, although the bilateral BLA and the direct BLA*→*NAc-S pathway were necessary to induce DOPr expression on CINs, and so to encode the specific S-O associations in the NAc-S, the BLA*→*NAc-S pathway was not engaged during their retrieval. Although optogenetic inhibition of BLA terminals in the NAc-S during Pavlovian conditioning blocked specific S-O encoding-related changes, the same inhibitory stimulation applied during the PIT test did not affect the animals’ ability to retrieve those associations: specific PIT was the same whether the BLA*→*NAc-S pathway was inhibited or not.

This finding forced us to consider other pathways through which the BLA might contribute to retrieval. Given previous evidence that the IL is involved in PIT (27), we began by comparing the BLA projection to the NAc-S with that to the IL and found little overlap between them (*≈* 3 %), with most BLA neurons projecting either to the NAc-S (*≈* 64 %) or the IL (*≈* 33 %). Furthermore, using an anterograde transsynaptic tracer to deliver Cre to the postsynaptic targets of the BLA in the IL, we established that these targets themselves project to the NAc-S, thereby completing a BLA*→*IL*→*NAc-S pathway. Next, we assessed the effect of optogenetically silencing the BLA*→*IL pathway on both the encoding and retrieval of the specific S-O associations that support PIT. Silencing this pathway blocked PIT whether applied during encoding or retrieval at test. We therefore hypothesized that the BLA*→*IL pathway engages the NAc-S during retrieval and tested this hypothesis using a novel disconnection approach in which the BLA*→*IL pathway was optogenetically silenced in one hemisphere and the IL*→*NAc-S pathway in the contralateral hemisphere during the PIT test. Relative to controls, specific PIT in the disconnection group was normal with the light off but was abolished when these pathways were inhibited with the light on, confirming that S-O retrieval on test required an intact BLA*→*IL*→*NAc-S pathway. This effect was anatomically specific: inhibiting the IL projection to the NAc-C did not affect PIT.

These data suggested that the encoding-related changes in the NAc-S might depend on the coordinated engagement of the IL inputs into the shell. However, lesioning the IL during S-O encoding did not affect DOPr expression on CINs in the NAc-S. Instead, it appears that both the BLA*→*NAc-S and BLA*→*IL pathways are necessary to encode the specific S-O associations that support PIT, but that only the latter pathway is required for their retrieval during the PIT test. Importantly, retrieval via the IL*→*NAc-S pathway depended on IL activation of NAc-S CINs. PIT was abolished when we optogenetically silenced the IL*→*NAc-S pathway in one hemisphere of ChAT-Cre rats in which CINs had been ablated into the contralateral NAc-S. These results demonstrate that IL modulation of local NAc-S CINs on test is required for the expression of outcome-specific PIT. Although we were unable to visualize or quantify DOPr expression on NAc-S CINs in the ChAT-Cre rats, the data are consistent with our broader account. In the CIN-ablated hemisphere, predictive learning was unable to establish the DOPr-based memory it normally generates, leaving only the contralateral hemisphere available to support transfer. Silencing the IL*→*NAc-S projection in that hemisphere removed the only remaining substrate through which the retrieval of predictive learning could influence choice, abolishing PIT. This interpretation aligns with our previous demonstration that DOPr accumulation in the NAc-S, while necessary for predictive learning to influence choice between actions, is not required for Pavlovian S-O learning per se (7).

Although the BLA*→*NAc-S and BLA*→*IL pathways differed in their involvement during the retrieval of specific S-O associations that support PIT, both were necessary for encoding those associations during initial Pavlovian conditioning. However, their involvement in that encoding differed. We found that the BLA input to the NAc-S is necessary to induce the specific changes in DOPr expression on CINs that underlie S-O encoding: inhibiting that input during exposure to one S-O association, but not during exposure to a second S-O association, abolished the effect of the former but not the latter on choice in the PIT test. For this pattern to emerge, the BLA input must convey specific S-O information to the NAc-S, modifying DOPr expression on distinct populations of CINs for different S-O associations. Interestingly, the accurate retrieval of these distinct associations appears to depend on the BLA inputs to the IL and, consequently, the IL inputs to NAc-S, pathway activity that must in some way be coordinated with the direct BLA inputs to the NAc-S.

Although these pathways appear to involve largely independent populations of BLA projection neurons, they differ in the time course over which they act. The modulatory processes engaged by the BLA*→*NAc-S input are relatively slow (4), whereas the BLA appears to engage pyramidal tract neurons in the IL rapidly (22). These IL neurons target distinct populations of spiny projection neurons (SPNs) in the NAc-S, one expressing dopamine D1 (D1-SPN) and the other D2 (D2-SPN) receptors (25). Whereas D1-SPNs appear to constitute the primary excitatory output for PIT, we suggest that the IL input to D2-SPNs may function to release enkephalin, which is expressed by these neurons and is the primary ligand for DOPr. This IL-induced release could, therefore, provide the conditions necessary to retrieve the BLA-induced GPCR-memory expressed on CINs. If transmission in the BLA*→*NAc-S and BLA*→*IL*→*NAc-S pathways is sufficiently temporally contiguous to allow changes in the BLA*→*NAc-S to engage with inputs from the IL, this may provide the plasticity necessary to target the appropriate D1-SPNs to capitalize, subsequently, on the excitatory drive provided by enkephalin-induced inhibition of CINs to generate the output for PIT.

How this D1-SPN activity mediates the bias in choice between instrumental actions has recently become clearer. Some time ago we found that PIT depends on the projection from the NAc-S to the medial ventral pallidum (mVP) (31), and we have recently shown that this output is mediated by D1-SPNs and not by D2-SPNs (30). From there, we found that the mVP generates inhibitory output to the mediodorsal thalamus (mdT), which inhibits the mdT (32). Recently, we found that this timed inhibition of the mdT acts to release mdT inhibition of the lateral orbitofrontal cortex, which increases its output to motor control processes in the dorsal striatum to induce PIT (33). Therefore, this research has uncovered a 521 broad but specialized circuit that allows Pavlovian conditioning to influence choice between instrumental actions but that plays no role in either Pavlovian or instrumental conditioning *per se*. The latter point was demonstrated repeatedly in the current experiments: although manipulations of the circuits engaged during the encoding and retrieval of S-O associations consistently influenced PIT, at no point did they affect the ability of outcome-devaluation to modify choice. Considerable evidence points to a distinct but parallel circuit mediating that latter process and involving the NAc-C and its connections to the BLA and insular cortex (34, 35), with output to the dorsomedial striatum via the medial prefrontal cortex (36) during encoding and the medial orbitofrontal cortex during retrieval (37). Therefore, these data support a sharp division in the circuitry mediating stimulus-based and value-based decision-making.

Beyond these broader conclusions about circuitry, the current experiments have important implications for theories of memory. On the basis of evidence from engram cells, the mechanisms of memory retrieval are often assumed to recapitulate those of encoding. Many memory functions, however, are heavily circuit based and can, therefore, engage different processes at encoding and retrieval. Here we found that, whereas both the BLA*→*NAc-S and BLA*→*IL circuits are involved in the encoding of specific stimulus-outcome associations, only the latter was required to retrieve those associations to bias choice between distinct instrumental actions during outcome-specific Pavlovian-instrumental transfer.

## MATERIALS AND METHODS

### Subjects

The subjects were experimentally naive DOPr-eGFP mice (20-35 g, 8-24 weeks; The Jackson Laboratory, stock number: 029012, strain code: B6;129S2-Oprd1tmKff/J) that express a fluorescent and functional form of DOPr, Long-Evans rats or ChAT-Cre rats (200-500 g, 8-24 weeks; breeding facility, UNSW) that express Cre recombinase in choline acetyltransferase neurons (ChAT). Although these efforts were limited by animal availability and postmortem analyses, we aimed to allocate an equal number of male and female animals in each group across all experiments. Note that our previous work found no influence of sex on PIT or value-based choice (38). All test data present individual performance for males and females. Animals were housed in transparent plastic boxes with their littermates (2-6 mice and 2-4 rats per box) throughout, and in a climate-controlled colony room maintained on a 12-hour light-dark cycle (lights on between 7:00 am and 7:00 pm). Water and standard lab chow were available *ad libitum* prior to the start of each experiment. Food restriction was initiated 3 days prior to the start of the protocols and was maintained throughout all training and testing phases. Animals were maintained at *≈* 90 % of their *ad libitum* body weight during the behavioral protocols and received a daily food amount that was adjusted based on their body weight measurements recorded every two days. The Animal Ethics Committee at the University of New South Wales approved all experimental procedures.

### Viruses

See Table 1 for a summary of viruses, surgical coordinates (indicated in mm from bregma) and animal strains used. The following viruses from AddGene (Watertown, MA, USA) were used: rAAV5/hSyn-hM4Di-mCherry (hM4Di; #50475), pAAV5/hSyn-mCherry (mCherry; #114472), AAV5-CaMKIIa-mCherry (mCherry; #114469) and AAV5/EF1a-DIO-eYFP (Cre-eYFP; #27056). The following viruses from UNC Vector Core (Chapel Hill, NC, USA) were used: pAAV5-hSyn-eYFP (eYFP; #117382), AAV5/hSyn-eNpHR3.0-eYFP-WPRE-PA (eNpHR; #26972), AAV5-CaMKIIa-eNpHR3.0-mCherry-WPRE (eNpHR; #4346), rAAV2/Ef1a-mCherry-IRES-WGA-Cre (WGA-Cre-mCherry; #55632) and AAV5/Flex-taCasp3-TEVP (Casp-Cre; #45580).

**Table 1.**
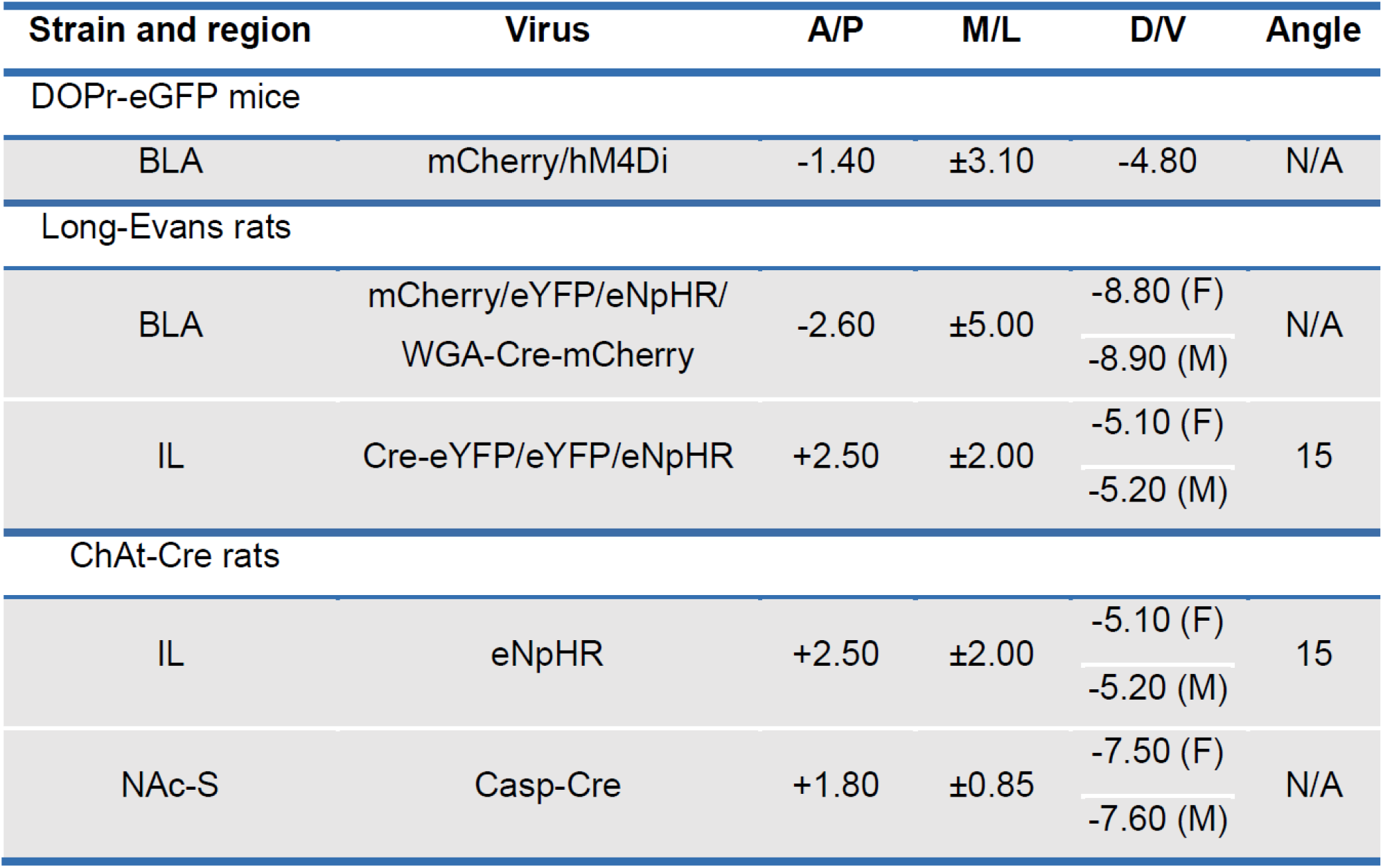
Summary of viruses, coordinates (indicated in mm from bregma) and strains used.

### Surgeries

Stereotaxic surgeries were performed under a continuous flow of isoflurane in oxygen (Mice: 0.6 L/min; induction 2-3 %, maintenance 1-1.5 %; Rats: 0.6 L/min; induction 4-5 %, maintenance 2-2.5 %). Animals were placed in a stereotaxic frame (Kopf Instruments, Tujunga, CA, USA), the surgical area was prepared with betadine iodine antiseptic solution, and a midline incision was made to expose the skull. At the incision site, bupivacaine hydrochloride (mice: 0.5 %; 0.1 mL; rats: 0.5 %; 0.2 mL) was injected subcutaneously, followed by subcutaneous administration of meloxicam (Metacam; 1 mg/kg) in the lower flank. The membrane over the skull was cleared, and the skull was levelled such that bregma and lambda were aligned in the same horizontal plane. For viral infusion, small holes were made in the skull above the target regions (mice: BLA ; rats: BLA, NAc-S and IL; see Table 1 for coordinates). In mice, viruses were infused using a pulled glass capillary (3.5”; Drummond Scientific) filled with mineral oil and mounted on a Nanoject III (Drummond Scientific). After a 2 min wait at the target site, 0.42 µL of virus was infused unilaterally at 23 nL/s, with 30 s intervals between pulses; the pipette was left in place for 3 min after the final pulse before withdrawal. In rats, infusions were performed using 1-µL glass syringes (86200; Hamilton Company, Reno, NV, USA) attached to an infusion pump (Standard Infuse/Withdraw Pump 11 Elite; Harvard Apparatus, Holliston, MA, USA), delivering 0.5 µL of virus unilaterally or bilaterally at 0.1 µL/min; the needle was left in place for 3 min following each infusion before being gently withdrawn. Following surgery, incisions were closed with staples (rats) or surgical silk sutures and sealed with tissue adhesive (mice; Vetbond, 3M), and the surgical area was treated with betadine iodine. Animals then received subcutaneous injections of the antibiotic Duplocillin (0.15 mL/kg) and sodium chloride (0.9 %; mice: 2 mL; rats: 4 mL) and were placed on a heating mat during recovery. They were returned to their home cages once alert and monitored daily, with *ad libitum* access to food and water, for a minimum recovery period of two to four weeks before behavioral testing.

Fiber-optic implantations in rats were conducted immediately after viral infusion, within the same surgery. Three or four jeweller’s screws were inserted into the skull, distributed anterior and posterior to the implantation sites. Fiber-optic cannulas (6 or 10 mm length, 0.48 mm diameter; Doric Lenses, Québec, QC, CAN) were then unilaterally or bilaterally implanted into the BLA, NAc-S or IL, and secured in place with dental cement (see Table 2 for coordinates).

**Table 2.** Summary of fiber-optic implantation coordinates (indicated in mm from bregma) and strains used.

| Strain and region | A/P | M/L | D/V | Angle |
| --- | --- | --- | --- | --- |
| Long-Evans rats |  |  |  |  |
| BLA | -2.60 | ±5.00 | -8.30 (F)<br>-8.40 (M) | N/A |
| IL | +2.50 | ±2.00 | -4.55 (F)<br>-4.70 (M) | 15 |
| NAc-S | +1.70 | ±2.19 | -6.85 (F)<br>-6.75 (M) | 10 |
| ChAt-Cre rats |  |  |  |  |
| NAc-S | +1.80 | ±0.85 | -6.85 (F)<br>-6.75 (M) | N/A |

The incision site was closed and secured with surgical silk sutures (68 601 4 G; Ethicon, North Ryde, NSW, AUS), and betadine antiseptic ointment was applied in the area.

### Drugs

The selective DOPr agonist SNC80 (Cayman Chemical, #156727) was dissolved in 0.1 M hydrochloric acid (HCl), and the pH was adjusted to 6-7 using 0.2 M sodium hydroxide (NaOH). It was then diluted in 0.9 % saline to reach a final concentration of 1 mg/mL. Mice received systemic intraperitoneal (i.p.) injections at 10 mg/kg. The vehicle solution was HCl and NaOH diluted in 0.9 % saline with an equivalent pH (7). The synthetic DREADD ligand Clozapine-N-Oxide (CNO; RTI International, #3423) was dissolved in distilled water and 0.1 % (v/v) 5 M HCl to reach a final concentration of 0.3 mg/mL. Mice received systemic i.p. injections at 3 mg/kg (39).

### Behavioral apparatus

Training and testing of mice and rats were conducted in sound- and light-attenuating operant chambers (MED Associates, Fairfax, VT, USA) controlled by MED-PC software. Chambers were illuminated by a 3 W, 24 V house light and contained a recessed feeding magazine centered on the right wall, equipped with an infrared photobeam to detect head entries. The magazine was connected to an individual food dispenser delivering grain pellets (Mice: 20 mg pellets, Bio-Serv #F0163; Rats: 45 mg pellets, Bio-Serv #F0165). Chambers were also fitted with a syringe pump (60 mL syringe) delivering 20 % sucrose solution into the magazine (Mice: 0.025 mL per delivery; Rats: 0.2 mL per delivery). Retractable levers were located on either side of the magazine. All chambers contained auditory stimulus devices, including a Sonalert white noise generator (80 dB), 3 kHz tone and 2 Hz mechanical clicker relay. Rat chambers (ENV-007-VP) were enclosed in light- and sound-resistant cabinets (ENV-018MD) and constructed with stainless-steel side walls, polycarbonate walls and ceiling, and a stainless-steel grid floor. Rat chamber ceilings included an access port for patch-cord connections in freely moving animals. Chamber activity was monitored via infrared cameras (Mice: D-Link DCS-932L; Rats: Dahua IPC-K35AP) using D-ViewCam software. All experimental events and behavioral responses (lever presses and magazine entries) were recorded automatically. Outcome-devaluation was conducted in a separate room equipped with 16 individual, open-top plastic boxes with stainless steel wire mesh lids. During devaluation, each chamber was fitted with either a ceramic dish for grain pellets or a plastic sipper bottle for sucrose solution.

### Optogenetic equipment

Fiber-optic cannulas consisted of a zirconia ferrule contained within a stainless-steel receptacle from which a silica/polymer flat-tip fiber extended. The patch cords (custom; Doric Lenses) were enclosed in a flexible cladding that consisted of one transparent fiber that split into two smaller fibers for attachment to the implanted cannulas.

Delivery of the LED light was controlled through a TTL adaptor which converted 28 V DC output to a TTL transition (MED Associates). The adaptor was connected to an LED driver (Doric Lenses) and connectorised LED light source (Doric Lenses) with a rotary joint for attachment to the patch cords. LED light was delivered as orange light (625 nm; continuous) and measured *≈* 10 mW at the cannula tip using a photometer (PM200; ThorLabs, Newton, NJ, USA) when the LED was switched on.

### Behavioral procedures

#### Pavlovian conditioning and retraining

Pavlovian conditioning involved eight 60 min daily sessions that consisted of presenting two stimuli (S1 and S2; tone, white noise or clicker), each paired with a particular food outcome (O1 and O2; grain pellets or sucrose solution; S1-O1 and S2-O2). Each stimulus was presented for 2 min, 4 times each in a pseudorandom order within each training session. Stimulus presentations were separated by pseudorandom intertrial intervals (ITIs) ranging from 3 to 5 min in length (4 min average). The stimulus order and ITIs were changed daily. Stimulus-outcome (S-O) pairings were counterbalanced within- and between-groups. Food outcomes were delivered on a random time 30 s schedule throughout the stimulus presentation (one outcome per delivery). Magazine entries were recorded throughout the session, and conditioned responses were analyzed by comparing magazine entries in the 2 min prior to stimulus presentation and during the 2 min stimulus presentation period. The additional Pavlovian conditioning session was identical to the initial training. For optogenetic manipulation in rats, when applicable, the LED was activated (625 nm; continuous) during each Pavlovian conditioning session, during either half of the stimulus presentation with their outcome or every presentation.

#### Instrumental conditioning

During instrumental conditioning, two actions (A1 and A2; left or right lever presses) were trained with the two different outcomes (O1 and O2) in 2 separate daily sessions for 10 days (mice) or 8 days (rats). The order of the sessions was counterbalanced, as were the action-outcome (A-O) relationships, which were further counterbalanced with the S-O relationships. Each session ended when 20 outcomes were earned or when 30 min had elapsed. For the first 4 days (mice) or 2 days (rats), actions were continuously reinforced (i.e. one action earned one outcome delivery). Over the next 3 sessions, actions were rewarded with a random ratio 5 (RR5) schedule (i.e. each action delivered an outcome with a probability of 0.2). For the final 3 days, animals were rewarded with an RR10 schedule (i.e. each action delivered an outcome with a probability of 0.1). Lever press actions were measured throughout the duration of the session, and performance during instrumental training was assessed using the mean number of lever presses per min.

#### Pavlovian-instrumental transfer test

During the test, both levers were inserted into the chamber simultaneously. The test was conducted in extinction (i.e., no outcomes were delivered). To reduce baseline lever presses, responding on both levers was extinguished for 8 min prior to any stimuli being presented. Each stimulus was presented for 2 min, 4 times each over the next 40 min in the following order: tone-noise-noise-tone-noise-tone-tone-noise (mice) or noise-clicker-clicker-noise-clicker-noise-noise-clicker (rats). Each stimulus presentation was separated by a 3 min fixed interval. For optogenetic manipulations in rats, when applicable, the LED was activated (625 nm; continuous) during half of the trials (on) but left inactive during the other half (off). The order for the on and off trials was counter-balanced. Lever press responses were recorded prior to and during stimulus presentations. These were then collated into “pre” (lever press rates in the 2 min prior to stimuli presentation), “same” (when the stimulus shares the same outcome with the action) and “different” (when the stimulus has a different outcome with the action) following data collection. In all experiments, “pre” was subtracted from “same” and “different” rates of responding to focus on the net effect of the predictive stimuli since baseline responding did not differ between groups across experiments.

#### Choice after outcome-devaluation test

Across the 2 consecutive days preceding these tests, rats were 691 given a daily instrumental conditioning session (RR10). Following each instrumental session, rats were habituated to the devaluation chambers for 30 min, where they had access to one outcome (counterbalanced). On the first test day, rats were placed in the devaluation chambers for 60 min where they had *ad libitum* access to one of the previously trained outcomes (O1 or O2). If optogenetic manipulations were conducted at test, once removed from the devaluation boxes, patch cords were attached to the fiber-optic cannulas and connected to the LED, and the first choice after outcome-devaluation test was conducted. Both levers were made available for 2 min each, 4 times during the session with an intertrial interval of 2 min where the levers were retracted. The LED (625 nm; continuous) was activated during 2 of the 2 min trials where the levers were available (on) and was left inactive during the other 2 trials (off). The order of the on and off trials was counterbalanced, as well as the devalued outcome. If no optogenetic manipulation occurred at tests, both levers were made available for a total of 5 min each. The following days, rats were given 2 instrumental conditioning sessions (RR10) to reinstate normal instrumental performance. The next day, a final choice after outcome-devaluation test was conducted as described above, except that the other outcome was devalued (counterbalanced). Lever presses and magazine entries were recorded while the levers were available.

### Tissue preparation and histology

#### Transcardial fixation and brain sectioning

Animals were rapidly euthanized with sodium pentobarbital i.p. injections and transcardially perfused with cold 4 % paraformaldehyde (PFA) in 0.1 M phosphate-buffered saline (PBS; pH 7.5). Brains were post-fixed in PFA at 4 °C overnight. Coronal sections (30 µm thick for mice and 40 µm for rats) were cut through the targeted structures with a vibratome (VT1200; Leica Microsystems, North Ryde, NSW, AUS) and stored at -20 °C in cryoprotective solution containing 30 % ethylene glycol, 30 % glycerol and 0.1 M PBS until they were processed for immunofluorescence.

#### Immunofluorescence

For DOPr and ChAT staining in mice, as well as ChAT staining and eGFP amplification in rats, individual free-floating sections were selected in 1X TBS (0.25 M Tris, 0.5 M NaCl, pH 7.5) to contain the region of interest and rinsed 3 times for 10 min in TBS solution. After a 2-hour incubation in blocking solution (0.5 % Triton X-100, 10 % NHS in TBS), sections were rinsed 3 times in TBS for 10 min each. Sections were then incubated between 12-48 hours (dependent on antibody) in a solution containing the primary antibodies (0.2 % Triton X-100, 2 % NHS in TBS) at 4°C. The primary antibodies were the following: polyclonal rabbit anti-eGFP (1:500; Thermo Fisher Scientific, #A-11122) and polyclonal goat anti-ChAT (1:300; Millipore, #AB144P). After the specified incubation period, the sections were washed 3 times in TBS for 10 min each and then transferred into a solution containing the secondary antibodies in TBS solution at room temperature for 1 hour. The secondary antibodies were the following: donkey anti-rabbit Alexa Fluor 488 (Figure 1C: 1:300; Figure 1H,N: 1:400; Thermo Fisher Scientific, #A-21206) and donkey anti-goat Cy3 (1:400; Jackson ImmunoResearch Laboratories, #705-165-147). Next, the sections were rinsed 3 times for 10 min in TBS and then mounted onto Superfrost Plus-coated slides using Vectashield hardset antifade mounting medium (H-1400; Vector Laboratories). Sections that did not require immunofluorescence staining were washed 3 times for 10 min each in TBS and mounted using Vectashield hardset antifade mounting medium.

### Microscopy and imaging

#### Fluorescence analysis and quantification

For DOPr imaging and quantification in mice, all images were obtained using sequential laser scanning microscopy (FV1000; Olympus). The somatic membrane distribution of DOPr was studied in individualized NAc-S CINs by obtaining a single confocal image of all ChAT immunoreactive neurons found in the region of interest (ROI; bilaterally) in tissue sections. An average of 15 neurons per hemisphere (a total of 2488 CINs were acquired in 95 mice for this study) were imaged using the confocal microscope with a 60 X objective (lens immersed in UPLSAPO 60 X oil). Optimal CINs were identified as those with a substantially intact membrane and a clearly visible nucleus. For each CIN, a single confocal plane was defined in the ChAT channel. Sequential 52.22 µm2 images (optical magnification: 60 X; digital zoom: 4 X, resolution; 19608 pixels/µm2) were obtained for ChAT signal (Ch02, HeNe green laser intensity is usually: 15 %; PMT: 680 V, offset: 5 %) and the corresponding eGFP signal (Ch01, Argon laser intensity is usually: 17 %; PMT: 750 V; offset: 40 %) with a Kalman filter (5 averaging scans). All fluorescent quantifications were performed on raw 16-bit images with ImageJ/Fiji software (40). We customized a workflow that measures DOPr fluorescence by writing an ImageJ/Fiji macro adapted from (17) and https://github.com/ahklemm/ImageJMacro_Introduction.

The workflow involves 8 steps: (1) First, we split the original multi-channel image stack into individual channels for independent processing. (2) Second, we blur the ChAT channel image to attenuate noise before (3) binarizing it by intensity thresholding. (4) Next, the “Analyze Particles” command is used as a filter to retrieve CIN outlines as well as removing small non-neuronal signals. (5) Using the previously retrieved outlines and the “Make Band” command, ROI 1 is defined on the somatic membrane (located in the intracellular-extracellular interface defined by the ChAT staining) of every CIN in the ChAT channel. (6) ROI 2 is defined with intensity thresholding and “Invert LUT” command, then used as a background correction and comprised the nuclear region (as defined by the central region devoid of ChAT staining). (7) Mean gray value for each ROI is collected from the overlapped eGFP image and expressed, for each neuron, as ROI 1 - ROI 2. (8) An average normalised DOPr fluorescence intensity is calculated for each mouse, and these values are averaged across groups.

To assess for viral infusion spread (mice and rats), as well as neuronal pathways and placement of fiber-optic cannulas (rats), imaging of the mounted brain sections was performed on a spinning disk confocal microscope (Zyla 4.2 sCMOS; Andor Technology, Belfast, UK) using the Diskovery multi-modal imaging platform (Oxford Instruments; Belfast, UK). The Diskovery platform was mounted on an inverted microscope body with a motorized stage and Perfect Focus System (Eclipse Ti; Nikon, Melville, NY, USA). Image capture was controlled using the Nikon NIS-Elements Software. Single high-magnification field-of-view images were generated by automatic stitching of multiple adjacent frames from within a defined area using the motorised stage (optical magnification: 20 X; pixel depth: 16-bit). Additional images were taken with a confocal microscope (BX61W1; Olympus, Shinjuku, TYO, JPN) using Fluoview software (FV1000; Olympus) for further placement analysis. All images were processed using ImageJ/Fiji software.

#### NAc-S CINs mapping

High-resolution mosaics of brain samples obtained with NIS-Elements were quantitatively analyzed using ImageJ/Fiji software. The outer boundary of the NAc-S was manually delineated in each hemisphere (freehand tool) in the eGFP channel, the area of the region was measured (mm2), and coordinates (x, y) of the points defining the line were obtained. To quantify the degree of NAc-S CIN ablation and retrieve the location of surviving CINs across the NAc-S, cell counts were conducted manually in the ChAT channel using ImageJ/Fiji software. Individual region of interest (ROI) was defined and coordinates (x, y) of each individual cell obtained. Cell counts were analyzed according to the total number of cells, calculated as the mean number of cells per mm2, per hemisphere, averaged across 1-2 slices per rat. Resulting neuronal data, including coordinates (x, y) and area for each detected ROI, were exported in a tab-delimited spreadsheet format and imported into MATLAB. A scatter plot with circles at the locations specified by the neuron’s coordinates was digitally overlaid onto a reconstructed outline of the corresponding NAc-S section (39).

#### BLA density mapping

To determine shared and independent projections from the BLA to IL and/or NAc-S, three sections of the BLA from anterior to posterior (-2.28, -2.50, -2.76 mm) in each animal were processed using ImageJ/Fiji software (1-2 sections per bregma coordinates per rat). The outer boundary of the BLA was manually delineated (freehand tool), the area of the region was measured (mm2), and coordinates (x, y) of the points defining the line were obtained. To quantify the number of mCherry and eGFP positive cells and retrieve their location across the BLA, images were separated per channel and automatically thresholded based on pixel intensity and individual nuclei were automatically detected using the “Analyze Particles” command in ImageJ/Fiji: it finds the edge of an object (i.e. nucleus), outlines its contour using the wand tool, creates an individual ROI, and then determines the coordinates (x, y) of its centre point. Resulting neuronal data, including position (x, y coordinates) and area of the eGFP and mCherry channel for each detected ROI, were exported in a tab-delimited spreadsheet format and imported into MATLAB. The position labelled neurons within the tissue were digitally reconstructed by combining a line plot defined by the edge points of the BLA and a scatter plot with circles at the locations specified by the neurons’ centroid coordinates. To visualize the density of labelled neurons targeting either the NAc-S, IL or both, we color coded single data points based on their calculated kernel density estimate (36, 39).

### Electrophysiological recordings

#### Brain slice preparation

Rats were euthanized under isoflurane gas anesthesia (4 % in air). Brains were immediately removed and sliced using a vibratome (VT1200; Leica Microsystems) in ice-cold oxygenated sucrose buffer solution (consisting of [in mM]: 241 sucrose, 28 NaHCO3, 11 glucose, 1.4 NaH2PO4, 3.3 KCl, 0.2 CaCl2 and 7 MgCl2). Coronal brain slices (300 µm thickness) containing the NAc were sampled and maintained at 33 °C in a submerged chamber containing physiological saline (consisting of [in mM]: 124 NaCl, 2.5 KCl, 1.25 NaH2PO4, 1 MgCl2, 1 CaCl2, 10 glucose and 26 NaHCO3) and equilibrated with 95 % O2 and 5 % CO2.

#### Recordings

Following 1 hour equilibration, slices were transferred to a recording chamber. An upright microscope (BX50WI; Olympus) was used to visualize neurons using differential interference contrast (DIC) Dodt tube optics and eYFP fluorescence, and slices were superfused continuously (1.5 mL/min) with oxygenated physiological saline at 33 °C. Whole-cell patch clamp recordings were performed using glass pipettes (tip size 2-5 MΩ) containing internal solution (consisting of [in mM]: 127 cesium methanesulfonate, 20 NaCl, 20 HEPES, 0.4 EGTA, 5 tetraethylammonium chloride, 3 QX314 chloride, 2.5 Mg-ATP, and 0.33 Na-GTP, pH 7.3, osmolarity 285-290 mOsm/L. Neurons sampled during recording were marked with Biocytin (0.1 %) added to the internal solution. A Multiclamp 700B amplifier (Molecular Devices; San Jose, CA, USA) connected to a Macintosh computer and interface ITC-18 (InstruTECH; Longmont, CO, USA) was used for data acquisition. Liquid junction potentials of - 10 mV were not corrected. Membrane currents were sampled at 5 kHz (low pass filter 2 kHz, Axograph X; Axograph, CA, USA) and voltage-clamped at - 80 mV. Series resistance (< 25 MOhm) was compensated by 75 % and monitored. Stock solutions of all drugs were diluted to working concentrations in the extracellular solution immediately before use and applied by continuous superfusion. Data from whole-cell recordings were included in analyses if the neurons appeared healthy under DIC on the monitor screen.

#### Electrical stimulation of BLA terminals

Evoked excitatory postsynaptic currents (EPSCs) were evoked with a pair of bipolar tungsten stimulating electrodes placed near (100-200 µm) the cell body, in the presence of picrotoxin (100 µM) to block GABAa receptors. Such stimulation most likely activates all axon terminals from various origins. Electrical stimulation (5 ms duration, using a Digitimer DS2A, UK) was applied at 20 s intervals. The intensity of stimulation was adjusted to obtain initial EPSC having an amplitude of 200-400 pA, such that both inhibition and augmentation could be observed with the superfusion of drugs or LED light. The stimulus intensity was not changed once the amplitude of the EPSC was steady. Simultaneous LED 625 nm illumination was achieved by a brief exposure of the recording neuron to a continuous wave LED (0.4 mW) for the duration of electrical pulses.

#### Post-hoc histological analysis

Immediately after physiological recording, brain slices containing biocytin-filled neurons were fixed overnight in 4 % PFA/0.16 M PB solution, washed and then transferred into 0.3 % Triton X-100/PB solution for 3 hours to permeabilize cells. Slices containing eYFP or eNpHR were then placed in 10 % NHS/PB for 1 hour before being incubated in primary chicken anti-GFP (1:1000; Aves Labs) for 2 hours at 4 °C to enhance fluorescent signal. The slices were rinsed in PB and then received a one-step incubation containing Alexa Fluor 488-conjugated donkey anti-chicken secondary antibody (1:500; Jackson Laboratories), and Cy3-conjugated avidin (1:1000; ExtrAvidin, Sigma) for 2 hours. Stained slices were rinsed in PB 3 times for 10 min each, mounted onto a glass slide, and then coverslipped with Fluoromount-G mounting medium (Southern Biotech, Birmingham, AL, USA). Neurons were imaged under a confocal microscope (Fluoview FV1000 and BX61WI, Olympus). Planned mixed contrasts were used to assess within-cell differences. A first within-subject contrast tested for differences between “baseline” and no LED light delivery (LED off). The second contrast tested for differences between these two conditions and LED illumination (LED on). The type-I error rate was set at 0.05 for each contrast tested.

### Statistical analyses

The data presented met the assumptions of the statistical tests used. The main measures were the number of magazine entries and lever presses per minute. The differences between groups were analyzed using a planned orthogonal contrast procedure 873 controlling the per contrast error rate. When applicable, a first contrast compared mice that had been injected with SNC80 (snc) or vehicle (veh), and rats that had been infused with either the control eYFP/mCherry virus or the inhibitory eNpHR virus. Changes across days during Pavlovian and instrumental conditioning were assessed by planned linear trend analyses. In mice, conditioned responding during the Pavlovian stage was analyzed using an elevation ratio of magazine entries, as this was the measure used in our previously published articles using mice (7, 17). This ratio was obtained by dividing the total number of magazine entries during S1 and S2 by the sum of that number with the total number of magazine entries in the pre-stimulus (pre) period [i.e., CS/(CS+pre)]. An elevation ratio of 0.5 indicated that the animals entered the magazine as much during the CS as outside the CS (i.e., poor learning). In contrast, an elevation ratio close to 1 showed that the animals entered the magazine substantially more in the presence of the CS than in its absence (i.e.,good learning). In rats, conditioned responding was analyzed by comparing magazine entries in the 2 min prior to stimulus presentation and during the 2 min stimulus presentation period. Pavlovian conditioning and the PIT tests included a within-subject factor comparing magazine entry rates during the 2 min stimuli against rates during a 2 min baseline period immediately preceding the stimuli. During the PIT tests, a within-subject factor examined the net (i.e., minus baseline) rates of lever presses on the response (same) earning the same outcome as predicted by the stimulus against those on the response (different) earning the different outcome. For optogenetic manipulations in rats, when applicable, a second within-subject factor compared performance when the LED was activated (on) or not (off). Value-based choice tests used a within-subject factor of response identity comparing rates of lever presses on the response (valued) earning the valued outcome against those on the response (devalued) earning the devalued outcome. When applicable, it also used the LED within-subject factor described above for rats that received optogenetic manipulations during the choice after outcome-devaluation test. All analyses were carried out using the PSY statistical software (School of Psychology, the University of New South Wales, Australia) or Prism 7 (GraphPad). The Type I error rate was controlled at alpha = 0.05 for each contrast tested. If interactions were detected, follow-up simple effects analyses were calculated to determine the source of the interactions. Data will be made available.

## ACKNOWLEDGEMENTS

Elise Pepin was supported by a Scientia PhD Scholarship from UNSW Sydney.

The research reported in this manuscript was supported by a Future Fellowship from the Australian Research Council (FT220100474) to V.L. and by a Senior Investigator Grant from the National Health and Medical Research Council of Australia (GNT1175420) to B.W.B.

## SUPPLEMENTARY MATERIALS

**Supplementary Figure S1.**
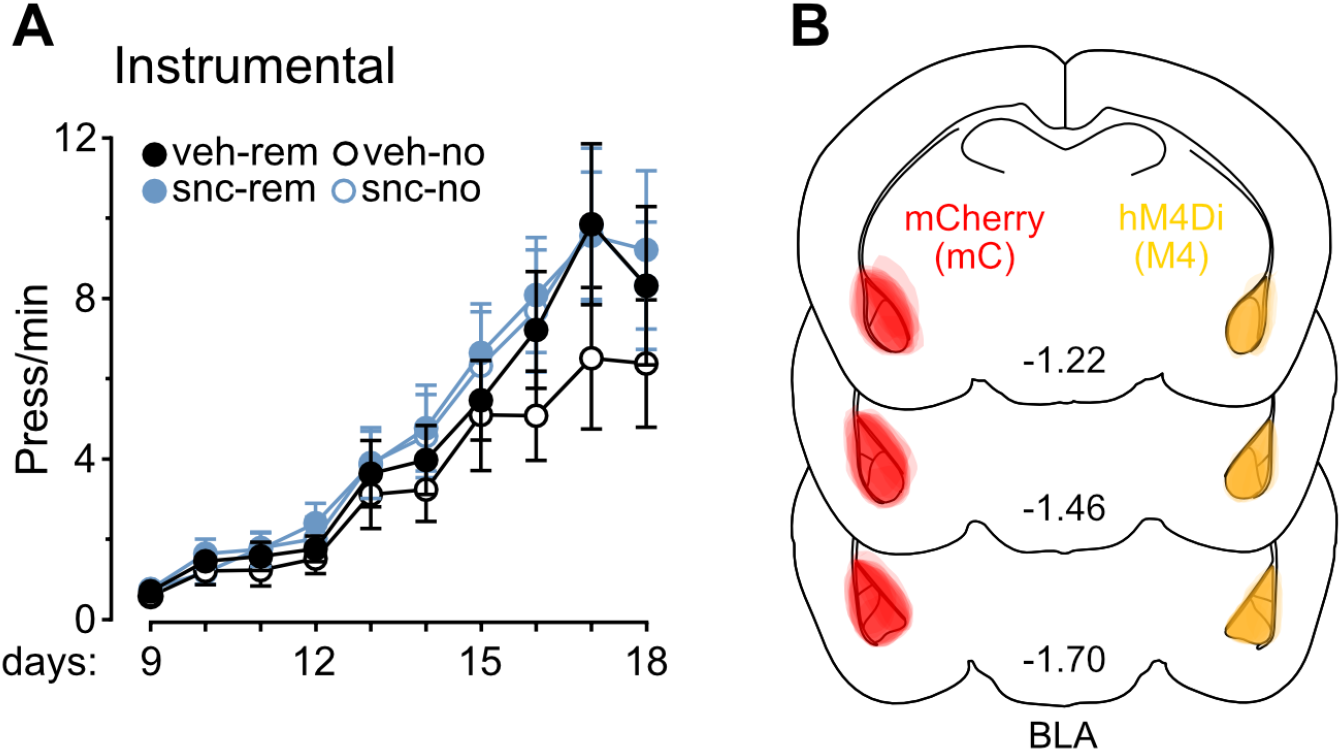
Histological and behavioral data related to Figure 1. **(A)** Instrumental training occurred smoothly, as mice displayed a gradual increase in lever press responding as training progressed (F(1,50)=82.75; p<0.001), regardless of groups (Fs<0.21). Overall lever press rates were similar across groups (Fs<0.2). **(B)** Representative micrographs showing the viral spread in the BLA of all mice. Darker regions represent the areas of greatest overlap across mice. Data are shown as mean ± SEM.

**Supplementary Figure S2.**
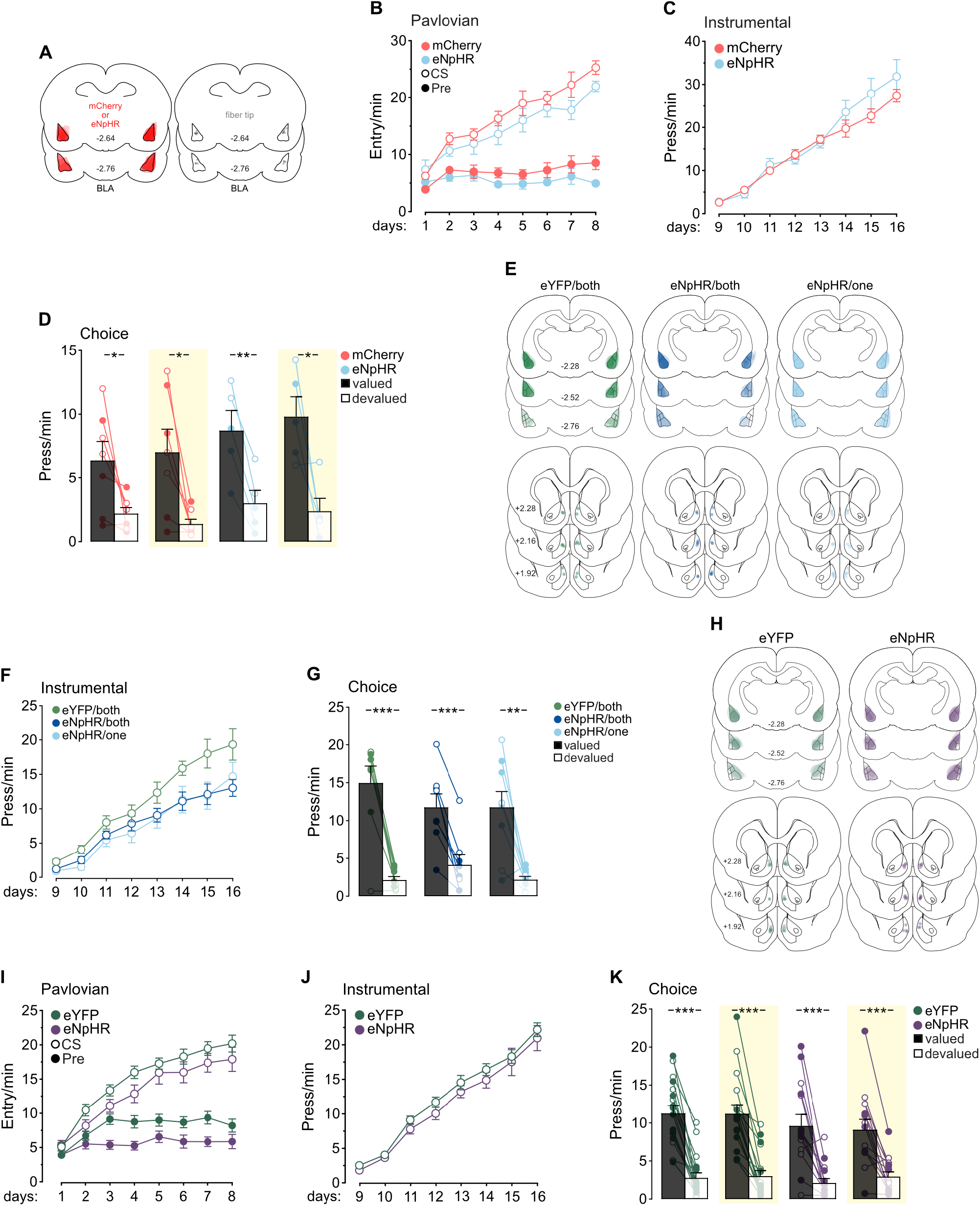
Histological and behavioral data related to Figure 2. **(A)** Data related to the BLA silencing at test experiment. Representative micrographs showing the viral spread (left) and fiber optic placements at their deepest position in the BLA of all rats (right). Darker regions represent the areas of greatest overlap across rats. **(B)** During Pavlovian conditioning, magazine entry rates were similar across groups (F<4.54) and increased over training (F(1,10)=206.30; p<0.001), with an interaction of day x group (F(1,10)=17.14; p<0.001). The rates were higher in the presence of the stimuli than in their absence (F(1,10)=111.07; p<0.001), irrespective of group (F<0.35). The discrimination between the two periods increased as training progressed (F(1,10)=77.70; p<0.001), regardless of group (F<0.34). **(C)** During instrumental conditioning, lever press rates were similar across groups (F<3.08) and increased as training progressed (F(1,10)=175.38; p<0.001), with a day x group interaction (F(1,10)=8.36; p<0.01). Although unexpected, this is unlikely to reflect any difference in learning per se, as evidenced by the presence of a devaluation effect in the choice test for both groups, under both light conditions. **(D)** During the choice after outcome-devaluation test, lever press rates were similar across groups (F<1.99) and were higher on the lever earning the valued outcome (F(1,10)=34.04; p<0.001) irrespective of group (F<0.72). Lever press rates were unaffected by LED activation (F<0.14) irrespective of group (F<0.07). The difference between rates on the valued and devalued lever was unaffected by LED activation (F<1.23) regardless of group (F<0.01). Simple effect analyses revealed that lever press rates were higher on the valued lever compared to the devalued lever in both groups whether the LED was off or on (mCherry off: F(1,6)=7.62; p<0.05; mCherry on: F(1,6)=10.55; p<0.05; eNpHR off: F(1,4)=36.59; p<0.01; eNpHR on: F(1,4)=10.65; p<0.05). Individual data points for females (open circles) and males (filled circles) are presented. Yellow rectangles represent LED light delivery at test. **(E)** Data related to the BLA*→*NAc-S silencing during Pavlovian training experiment. Representative micrographs showing the viral spread in the BLA (top), and optic fiber placements at their deepest position in the NAc-S of all rats (bottom). Darker regions represent the areas of greatest overlap across rats. **(F)** During instrumental training, overall lever press rates were higher in the eYFP group relative to the eNpHR groups (eYFP/both vs. eNpHR/both + eNpHR/one: F(1,22)=6.74; p<0.05), with no difference between eNpHR groups (eNpHR/both vs. eNpHR/one: F<0.03). Rats displayed a gradual increase in lever press responding as training progressed (F(1,22)=177.42; p<0.001), with no day x group interaction (Fs<4.19). **(G)** During the choice after outcome-devaluation test, lever press rates were similar between groups (Fs<0.51) and were higher on the lever earning the valued outcome (F(1,22)=80.51; p<0.001) irrespective of group (Fs<3.11). Simple effect analyses revealed that lever press rates were higher on the valued lever compared to the devalued lever in all groups (eYFP/both: F(1,7)=36.98; p<0.001; eNpHR/both: F(1,7)=36.84; p<0.001; eNpHR/one: F(1,8)=19.17; p<0.01 ). Individual data points for females (open circles) and males (filled circles) are presented. **(H)** Data related to the BLA*→*NAc-S silencing at test experiment. Representative micrographs showing the viral spread in the BLA (top), and fiber optic placements at their deepest position in the NAc-S of all rats (bottom). Darker regions represent the areas of greatest overlap across rats. **(I)** During Pavlovian conditioning, eYFP rats showed higher magazine entry rates relative to the eNpHR group (F(1,29)=5.12; p<0.05). These rates increased over training (F(1,29)=110.06; p<0.001), regardless of group (F<0.19). Rates were higher in the presence of the stimuli than in their absence (F(1,29)=250.46; p<0.001), irrespective of group (F<0.30). The discrimination between the two periods increased as training progressed (F(1,29)=168.02; p<0.001), regardless of group (F<0.23). **(J)** Lever press responding increased over instrumental training (F(1,29)=375.02; p<0.001), regardless of group (F<0.06). Overall lever press rates were similar across groups (F<1.21). **(K)** During the choice after outcome-devaluation test, lever press rates were similar across groups (F<1.07) and were higher on the lever earning the valued outcome (F(1,29)=185.19; p<0.001) irrespective of group (F<1.86). Lever press rates were unaffected by LED activation (F<0.07) irrespective of group (F<0.01). The difference between rates on the valued and devalued lever was unaffected by LED activation (F<0.49) regardless of group (F<0.22). Simple effect analyses revealed that lever press rates were higher on the valued lever compared to the devalued lever in both groups whether the LED was off or on (eYFP off: F(1,16)=91.09; p<0.001; eYFP on: F(1,16)=71.05; p<0.001; eNpHR off: F(1,13)=23.52; p<0.001; eNpHR on: F(1,13)=30.21; p<0.001). Individual data points for females (open circles) and males (filled circles) are presented. Yellow rectangles represent LED light delivery at test. Data are shown as mean ± SEM. Asterisks denote significant effects (*: p<0.05; **: p<0.01; ***:p<0.001).

**Supplementary Figure S3.**
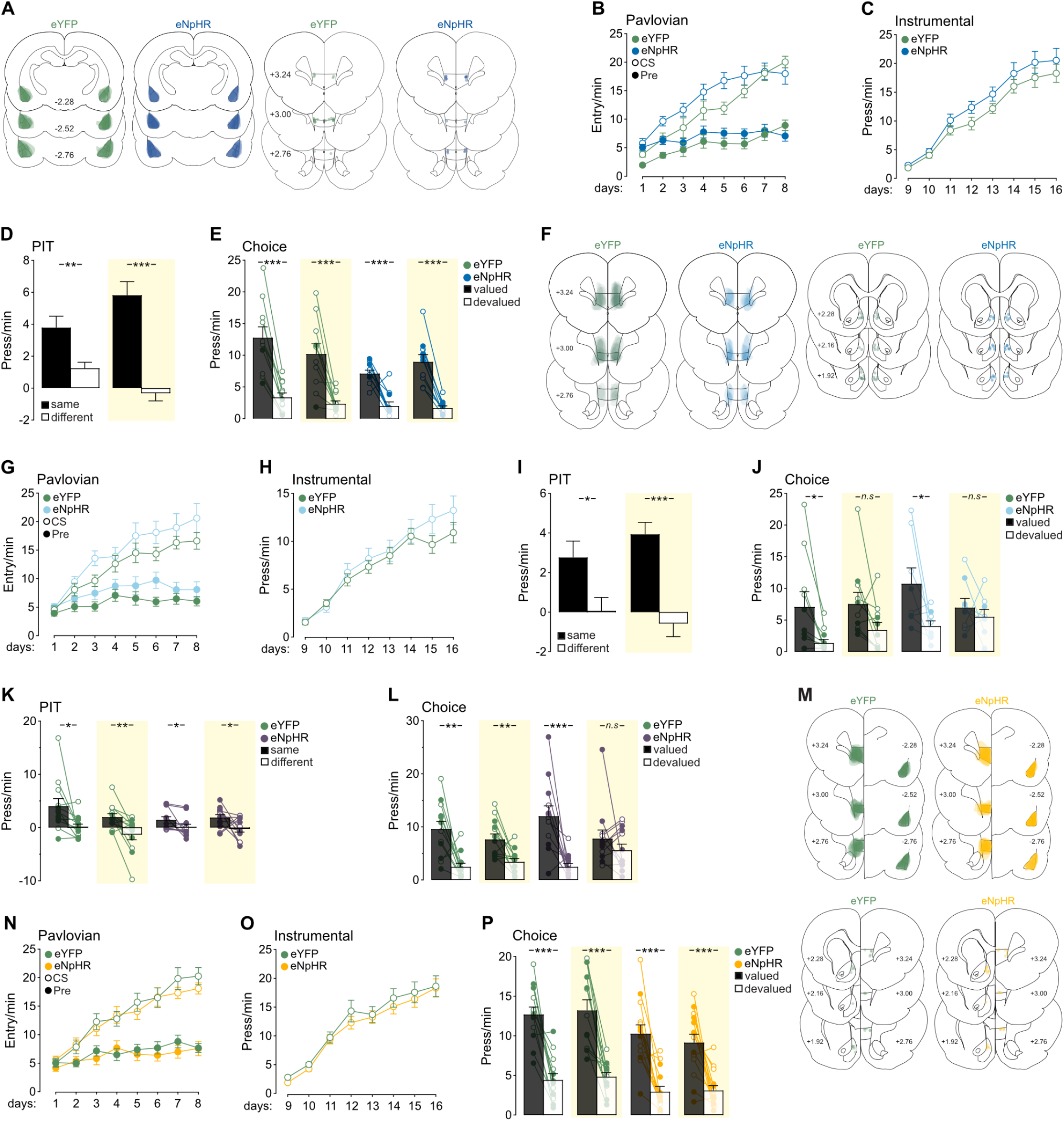
Histological and behavioral data related to Figure 3. **(A)** Data related to the experiments assessing BLA*→*IL silencing at test. Representative micrographs showing the viral spread in the BLA (left) and fiber optic placement at their deepest position in the IL of all rats (right). Darker regions represent the areas of greatest overlap across rats. **(B)** During Pavlovian conditioning, magazine entry rates were similar across groups (F<2.47) and increased over training (F(1,19)=160.02; p<0.001). The increase was more pronounced in the eNpHR group than in the eYFP group (F(1,19)=5.95; p<0.05). Rates were higher in the presence of the stimuli than in their absence (F(1,19)=264.82; p<0.001) irrespective of group (F<0.86). The discrimination between the two periods increased as training progressed (F(1,19)=118.70; p<0.001) regardless of group (F<0.01). **(C)** Lever press responding increased over instrumental training (F(1,19)=190.66; p<0.001) regardless of group (F<0.58). Overall lever press rates were similar across groups (F<2.11). **(D)** Unilateral silencing of the BLA*→*IL pathway left outcome-specific PIT unaffected in eNpHR animals that were excluded due to viral or fiber optic misplacement, leaving one hemisphere intact (excluded eNpHR rats from the BLA*→*IL pathway silencing at test in Figures 3I-L, and BLA*→*IL and IL*→*NAc-S pathways disconnection in Figures 3Q-T; *n*=16; 11 M, 5 F). Lever press rates during the test were similar between light conditions (F<0.22) and higher on the action earning the same outcome as the stimuli compared to the action earning the different outcome (F(1,15)=49.99; p<0.001), with a light x lever interaction (F(1,15)=22.78; p<0.001). Simple effect analyses revealed that lever press rates were higher on the same lever compared to the different lever whether the LED was off (F(1,15)=11.35; p<0.01) or on (F(1,15)=80.64; p<0.001). Yellow rectangle represents LED light delivery at test. **(E)** During the choice test, lever press rates were greater for eYFP rats relative to eNpHR rats (F(1,19)=4.43; p<0.05). These rates were higher on the lever earning the valued outcome (F(1,19)=94.20; p<0.001) irrespective of group (F<2.54). Lever press rates were unaffected by LED activation (F<0.91), but there was an interaction of light x group (F(1,19)=5.61; p<0.05). The difference between rates on the valued and devalued lever was unaffected by LED activation (F<0.09) regardless of group (F<3.51). Simple effect analyses revealed that lever press rates were higher on the valued lever compared to the devalued lever in both groups whether the LED was off or on (eYFP off: F(1,10)=54.36; p<0.001; eYFP on: F(1,10)=21.84; p<0.001; eNpHR off: F(1,9)=23.73; p<0.001; eNpHR on: F(1,9)=65.09; p<0.001). Individual data points for females (open circles) and males (filled circles) are presented. Yellow rectangles represent LED light delivery at test. **(F)** Data related to the IL*→*NAc-S silencing at test experiment. Representative micrographs showing the viral spread in the IL (left) and fiber optic placement at their deepest position in the NAc-S of all rats (right). Darker regions represent the areas of greatest overlap across rats. **(G)** During Pavlovian conditioning, magazine entry rates were similar across groups (F<2.49) and increased over training (F(1,28)=77.41; p<0.001) regardless of group (F<1.09). Rates were higher in the presence of the stimuli than in their absence (F(1,28)=151.33; p<0.001) irrespective of group (F<0.23). The discrimination between the two periods increased as training progressed (F(1,28)=130.96; p<0.001), regardless of group (F<0.91). **(H)** Lever press responding increased over instrumental training (F(1,28)=142.55; p<0.001), regardless of group (F<1.96). Overall lever press rates were similar across groups (F<0.79). **(I)** Unilateral silencing of the IL*→*NAc-S pathway left outcome-specific PIT unaffected in eNpHR animals that were excluded due to viral or fiber optic misplacement, leaving one hemisphere intact (excluded eNpHR rats from the IL*→*NAc-S pathway silencing at test in Figures 3M-P, and BLA*→*IL and IL*→*NAc-S pathway disconnection in Figures 3Q-T; *n*=12; 6 M, 6 F). Lever press rates during the test were similar between light conditions (F<0.24) and higher on the action earning the same outcome as the stimuli compared to the action earning the different outcome (F(1,11)=13.09; p<0.01), with no light x lever interaction (F<3.36). Simple effect analyses revealed that lever press rates were higher on the same lever compared to the different lever whether the LED was off (F(1,11)=5.44; p<0.05) or on (F(1,11)=18.01; p<0.001). Yellow rectangles represent LED light delivery at test. **(J)** A subset of rats in each group (eYFP: *n*=10, 5 M, 5 F; eNpHR: *n*=8, 4 M, 4 F) were given value-based choice tests. Lever press rates were similar across groups (F<1.50) and were higher on the lever earning the valued outcome (F(1,16)=14.26; p<0.01) irrespective of group (F<0.12). These rates were unaffected by LED activation (F<0.01) irrespective of group (F<2.36). The difference between rates on the valued and devalued lever was unaffected by LED activation (F<3.02) regardless of group (F<0.85). Simple effect analyses revealed that lever press rates were higher on the valued lever compared to the devalued lever, but this depended on the LED conditions in both groups (eYFP off: F(1,9)=8.25; p<0.05; eYFP on: F<4.28; eNpHR off: F(1,7)=5.96; p<0.05; eNpHR on: F<0.52). Individual data points for females (open circles) and males (filled circles) are presented. Yellow rectangles represent LED light delivery at test. **(K)** Additional experiment examining choice after outcome-devaluation in Figure S3J when silencing the IL*→*NAc-S pathway. To provide circuit specificity, the following experiment examined whether the IL*→*NAc-C pathway is necessary for outcome-specific PIT and choice tests. Using the same behavioural paradigms and optogenetics parameters as depicted above, two groups of Long-Evans rats were used: eYFP (*n*=12; 7 M, 5 F) and eNpHR (*n*=12; 7 M, 5 F). Silencing the IL*→*NAc-C pathway had no effect when it was conducted at the time of outcome-specific PIT test. Baseline responding (F<0.81) and overall lever press rates during the test were similar between groups (F<0.52) and light conditions (F<1.67), with no group x light interaction (F<1.89). Lever press rates were higher on the action earning the same outcome as the stimuli compared to the action earning the different outcome (F(1,22)=16.64; p<0.001), regardless of group (F<1.96). There was no light x lever interaction (F<0.01), and no light x lever x group interaction during the presentation of the predictive stimuli (F<0.90). Simple effect analyses revealed that lever press rates were higher on the same lever compared to the different lever in both groups whether the LED was off or on (eYFP off: F(1,11)=6.80; p<0.05; eYFP on: F(1,11)=11.95; p<0.01; eNpHR off: F(1,11)=6.57; p<0.05; eNpHR on: F(1,11)=4.91; p<0.05). Individual data points for females (open circles) and males (filled circles) are presented. Yellow rectangles represent LED light delivery at test. **(L)** Silencing the IL*→*NAc-C pathway abolished value-based choice. Lever press rates were similar across groups (F<1.41) and were higher on the lever earning the valued outcome (F(1,22)=28.24; p<0.01) irrespective of group (F<0.01). These rates were unaffected by LED activation (F<0.90) irrespective of group (F<0.01). The difference between rates on the valued and devalued lever was affected by LED activation (F(1,22)=10.62; p<0.01) but generated no light x group interaction (F<2.01). Simple effect analyses revealed that lever press rates were higher on the valued lever compared to the devalued lever, but this depended on the LED conditions in the eNpHR group (eYFP off: F(1,11)=14.56; p<0.01; eYFP on: F(1,11)=11.12; p<0.01; eNpHR off: F(1,11)=18.44; p<0.001; eNpHR on: F<1.01). Individual data points for females (open circles) and males (filled circles) are presented. Yellow rectangles represent LED light delivery at test.**(M)** Data related to the BLA*→*IL and IL*→*NAc-S pathways silencing at test experiment. Representative micrographs showing the viral spread in the IL and the BLA (top), as well as fiber optic placement at their deepest position in the IL and NAc-S of all rats (bottom). Darker regions represent the areas of greatest overlap across rats. **(N)** During Pavlovian conditioning, magazine entry rates were similar across groups (F<0.29) and increased over training (F(1,25)=100.08; p<0.001) regardless of group (F<0.97). Rates were higher in the presence of the stimuli than in their absence (F(1,25)=158.20; p<0.001) irrespective of group (F<0.01). The discrimination between the two periods increased as training progressed (F(1,25)=154.85; p<0.001) regardless of group (F<1.46). **(O)** Lever press responding increased over instrumental training (F(1,25)=172.18; p<0.001) regardless of group (F<0.01). Overall lever press rates were similar across groups (F<0.52). **(P)** During the choice test, lever press rates were greater for eYFP rats relative to eNpHR rats (F(1,25)=8.88; p<0.01). These rates were higher on the lever earning the valued outcome (F(1,25)=100.80; p<0.001), irrespective of group (F<1.18). Lever press rates were unaffected by LED activation (F<0.01) irrespective of group (F<1.27). The difference between rates on the valued and devalued lever was unaffected by LED activation (F<0.28) regardless of group (F<0.40). Simple effect analyses revealed that lever press rates were higher on the valued lever compared to the devalued lever in both groups whether the LED was off or on (eYFP off: F(1,12)=44.78; p<0.001; eYFP on: F(1,12)=36.34; p<0.001; eNpHR off: F(1,13)=29.72; p<0.001; eNpHR on: F(1,13)=21.15; p<0.001). Individual data points for females (open circles) and males (filled circles) are presented. Yellow rectangles represent LED light delivery at test. Data are shown as mean ± SEM. Asterisks denote significant effects (*: p<0.05; **: p<0.01; ***: p<0.001; *n*.*s*.: non-significant).

**Supplementary Figure S4.**
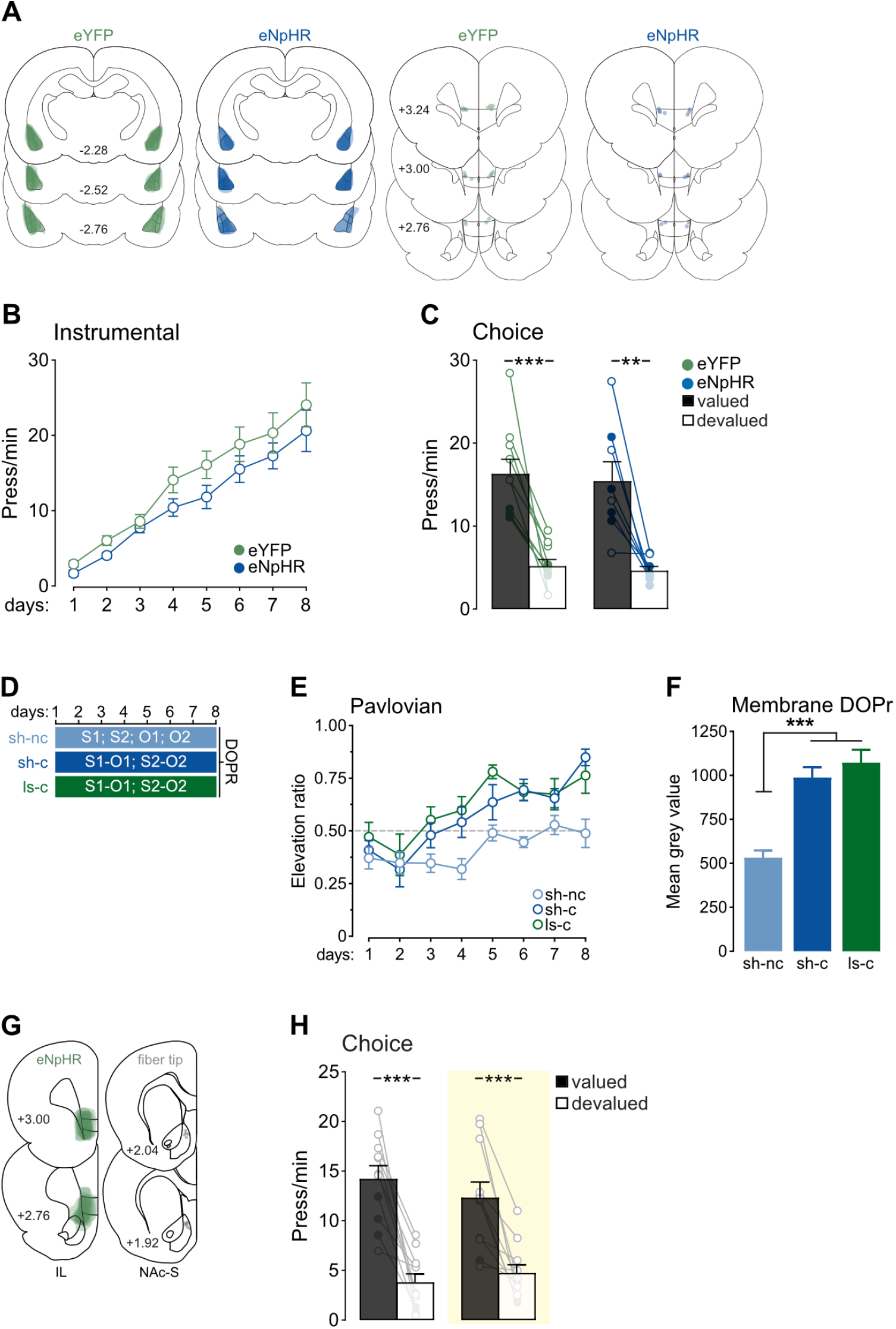
Histological and behavioral data related to Figure 4. **(A)** Data related to the BLA*→*IL silencing during Pavlovian training experiment. Representative micrographs showing the viral spread in the BLA (left) and fiber optic placement at their deepest position in the IL of all rats (right). Darker regions represent the areas of greatest overlap across rats. **(B)** Lever press responding increased over instrumental training (F(1,16)=101.54; p<0.001) regardless of group (F<0.37). Overall lever press rates were similar across groups (F<1.91). **(C)** During the choice test, lever press rates were similar between groups (F<0.22) and were higher on the lever earning the valued outcome (F(1,16)=50.41; p<0.001) irrespective of group (F<0.01). Simple effect analyses revealed that lever press rates were higher on the valued lever compared to the devalued lever in both groups (eYFP: F(1,9)=31.33; p<0.001; eNpHR: F(1,7)=20.25; p<0.01). Individual data points for females (open circles) and males (filled circles) are presented. **(D)** Additional experiment evaluating the role of the IL in the accumulation of DOPr on NAc-S CINs during Pavlovian conditioning. Three groups of DOPr-eGFP mice were used: sh-nc (*n*=8; 4 M, 4 F); sh-c (*n*=8, 4 M, 4 F) and ls-c (*n*=8, 4 M, 4 F). Mice received bilateral IL injections of either sham (sh-nc and sh-c) or ibotenic acid to induce a lesion (ls-c). After recovery, two groups received contingent (sh-c and ls-c) Pavlovian training. The non-contingent group (sh-nc) received the same number of stimuli and outcomes than sh-c and ls-c groups, although presentation of stimuli did not predict the delivery of the outcomes. Immediately after training, we quantified DOPr expression on NAc-S CINs of all mice. **(E)** Animals in the sh-c and ls-c groups learned that the two stimuli predicted the two food outcomes. Baseline responding was equivalent in all groups (F<2.2). Sham or ibotenic acid-induced lesions of the IL had no effect on conditioned responding: mice entered the magazine more during the stimuli than in their absence, as indicated by an elevation ratio above 0.5. This elevation ratio was similar between groups (F<0.7) and gradually increased across training (F(1,21)=63.36; p<0.001) irrespective of drug treatment (F<1.5). In contrast, mice receiving non-contingent training did not show evidence of predictive learning, as revealed by an elevation ratio of approximately 0.5. **(F)** Membrane DOPr expression was similar between contingently trained mice whether they had received excitotoxic IL lesion or not (sh-c vs. ls-c; F<0.7). Critically, this lack of difference was due to contingent training, as mice receiving non-contingent Pavlovian training showed lower DOPr expression than all other groups (sh-c + ls-c vs. sh-nc; F(1,21)=46.21; p<0.001). **(G)** Data related to the IL*→*NAc-S silencing at test experiment combined with contralateral NAc-S CINs lesion. Representative micrographs showing the viral spread in the IL (left) and fiber optic placement at their deepest position in the NAc-S of all rats (right). Darker regions represent the areas of greatest overlap across rats. **(H)** During the choice test, lever press rates were higher on the lever earning the valued outcome (F(1,10)=62.70; p<0.001) and unaffected by LED activation (F<0.38). The difference between rates on the valued and devalued levers was unaffected by LED activation (F<1.57). Simple effect analyses revealed that lever press rates were higher on the valued lever compared to the devalued lever whether the LED was off or on (off: F(1,10)=42.47; p<0.001; on: F(1,10)=22.26; p<0.001). Individual data points for female (open circle) and male (filled circle) are presented. The yellow rectangle represents LED light delivery at test. Data are shown as mean ± SEM. Asterisks denote significant effects (**: p<0.01; ***: p<0.001).

## Notes

### Competing Interest Statement

The authors have declared no competing interest.

